# Integrated transcriptomic analysis reveals a metabolically quiescent state and gene expression networks related to intermediate and mature human 8-cell stage embryo resembling cells *in vitro*

**DOI:** 10.1101/2025.09.03.673953

**Authors:** Pauliina Paloviita, Sonja Nykänen, Sanna Vuoristo

## Abstract

Human early development is challenging to study due to limited samples and cell numbers. The emergence of 8-cell stage embryo-like cells (8CLCs) offers new opportunities to understand embryonic genome activation (EGA) in humans. Our research compares and characterizes 8CLCs from various stem cell-based systems to determine how well these models reflect human early embryonic development. Using single-cell RNA sequencing (scRNA-seq) datasets from multiple studies, we integrated data to identify key gene co-expression modules, transposable element (TE) expression, and biological processes recapitulated in 8CLCs. We identified both mature and intermediate 8CLCs, with the Yoshihara and Mazid datasets best representing 8-cell stage embryos. 8CLCs show quiescence in energy and RNA metabolism, regulation of RNA splicing, and ribosome biogenesis, mirroring human 8-cell stage embryos. Our findings underscore the importance of distinguishing mature 8CLCs from partially reprogrammed cell states to improve their use as models for human EGA, *in vitro*.

## Introduction

The onset of transcription from the embryonic genome, embryo genome activation, demarcates the transfer of development guidance from maternal to embryonic transcripts. Human EGA occurs in two waves at the 4- to 8-cell stages (Töhönen et al., 2015) and failure to launch EGA results in defects in pre-implantation embryos (Niakan et al., 2012). While animal models have been useful in deciphering human developmental processes, these studies are hindered by interspecies differences as mice lack for instance EGA transcript *LEUTX* (Jouhilahti et al., 2016). Human EGA is technically and ethically challenging to study, due to the limited numbers of donated samples and inability to follow up development. Despite EGA being crucial for human reproduction and health, the molecular underpinnings remain largely understudied due to previous lack of sufficient *in vitro* models.

Recently, *in vitro* models of the human 8-cell stage embryos called 8-cell-like cells (8CLCs) were discovered (Balaton & Pasque, 2022; Mazid et al., 2022; Moya-Jódar et al., 2023; Taubenschmid-Stowers et al., 2022; Yoshihara et al., 2022; X. Yu et al., 2022). Spontaneously emerging 8CLCs exist in various human naïve pluripotent stem cell (hPSCs) backgrounds that have been cultured under distinct naive conditions, such as 5iLA and PXGL (G. Guo et al., 2017; Moya-Jódar et al., 2023; Theunissen & Jaenisch, 2017), *in vitro.* 8CLCs make up about 1-2% of naive hPSCs cultures, similar to the minor population of the mouse equivalent, 2-cell like cells (2CLCs), in mouse embryonic stem cells (mESCs) (Macfarlan et al., 2012; Rodriguez-Terrones et al., 2018). 8CLCs can also be induced genetically, for instance using DUX4 induction in naïve or primed hESCs (Taubenschmid-Stowers et al., 2022; Yoshihara et al., 2022), or chemically (Mazid et al., 2022; X. Yu et al., 2022) using histone deacetylase inhibition, histone H3K27me3 methyltransferase (EZH2) inhibition, PARP1 inhibition, and p53 activation among others (summarized in Taubenschmid-Stowers & Reik, 2023 and Yoshihara & Kere, 2023). These studies reveal that naive hPSCs have an intrinsic capacity to generate a transient 8-cell-like state, akin to 2CLCs in mice, with circulatory transitions inferred between naive hPSCs and 8CLCs (Taubenschmid-Stowers & Reik, 2023). While DUX4 induction or optimized medium composition can enhance the generation of 8CLCs, this effect is also temporary. 8CLCs have been sustained in cell death inhibitory culture conditions for up to 2-3 weeks (X. Yu et al., 2022) and recently a pre-EGA-like state was generated and stabilized via splicing inhibition using naïve and primed hPSC as starting material (Li et al., 2024), indicating that long-term culture of 8CLCs may be challenging but possible.

Single-cell RNA sequencing (scRNA-seq) with downstream analyses, including dimensionality reduction, clustering, and observation of 8-cell stage marker gene expression such as *ZSCAN4*, *TPRX1*, and *LEUTX*, as well as algorithmic cell annotation and integration with human embryo datasets, have been used to identify 8CLCs. Differences between initial reports and re-analysis of 8CLCs in naive hESC cultures have been noted, which may be attributable to varying approaches used to identify 8CLCs from scRNA-seq data (Yoshihara & Kere, 2023; X. Yu et al., 2022). For instance, in a previous integrative comparison using pre-processed scRNA-seq data of 8CLCs, Yoshihara & Kere found that many cells annotated as 8CLCs more closely resembled later embryonic stages than 8-cell stage embryos (Yoshihara & Kere, 2023). While biological differences likely exist between reported 8CLCs, the outcome of comparisons or subsequent validation efforts may depend on the stringency of initial criteria used to distinguish 8CLCs from other cells, including partially EGA-resembling cells as reported in mESCs (Rodriguez-Terrones et al., 2018).

Furthermore, not only gene expression patterns but also the expression of specific TEs demarcate the 8-cell stage in human embryos (Göke et al., 2015; Grow et al., 2015; Hashimoto et al., 2021; Liu et al., 2019; Pontis et al., 2019; Xu et al., 2022). Transcription of TEs, such as MERVL retrotransposons, is required for mouse preimplantation embryo development (Sakashita et al., 2023). Several studies have suggested that TE profiles of 8CLCs resemble those of 8-cell stage embryos by examining the expression or chromatin openness of selected TEs, such as MLT2A1 and MLT2A2 (Göke et al., 2015; Hashimoto et al., 2021; Liu et al., 2019; Mazid et al., 2022; Moya-Jódar et al., 2023; Taubenschmid-Stowers et al., 2022). However, a comparison of TE-mediated regulation between spontaneous, DUX4-induced, and medium-induced 8CLCs has not been conducted, which is essential since DUX4 is known to activate TEs (Geng et al., 2012; Hendrickson et al., 2017; Vuoristo et al., 2022; Whiddon et al., 2017).

While the regulation of EGA has been actively studied for several decades, it remains only partly understood, and it is unclear to what extent these findings apply to the molecular roadmap for generating 8-cell-like cells (8CLCs). Recently, Zou et al. (2022) discovered that in human embryos, PRD-like homeobox transcription factors *TPRX1/2/L* act upstream of other human EGA genes, including 8CLC markers *ZSCAN4* and *DUXA*. Additionally, TPRX1 and the methylation regulator DPPA3 were functionally shown to promote the upregulation of EGA gene expression and the generation of 8CLCs (Mazid et al., 2022). The binding motifs of PRD-like transcription factors DUX4, DUXA, and EGA-upregulated KLF17 were reported as enriched around 8CLC signature genes, suggesting their relevance in the spontaneous generation of 8CLCs (Taubenschmid-Stowers et al., 2022). While several evolutionarily related PRD and PRD-like homeobox transcription factors are very likely upstream regulators of 8CLC marker genes (Madissoon et al., 2016), it remains unclear which factors are indispensable in this process. Moreover, understanding whether EGA factors are differentially regulated in DUX4-induced, culture medium-induced, or spontaneously emerging 8CLCs, as well as identifying the common molecular hurdles that block 8CLC generation from hPSCs, provides a crucial basis for refining current 8CLC-related methods and ensures their adequate use to model human EGA *in vitro*.

## RESULTS AND DISCUSSION

### The expression of EGA genes distinguishes between intermediate and mature 8CLC identities

The human embryo research field has long sought an *in vitro* model for studying EGA, and recent publications introducing various 8CLC models necessitate comparison to understand their limitations and implications. To compare various 8CLC models, we identified optimal conditions for generating 8CLCs from original publications and retrieved raw reads for the scRNA-seq datasets (Fig 1A, Table S1). For spontaneously emerging 8CLCs, which constitute under 2% of naïve hESC cultures, no varying conditions were tested, and the TS (Taubenschmid-Stowers et al., 2022) and MJ (Moya-Jódar et al., 2023) datasets were used as provided. For Mazid (Mazid et al., 2022), we used data from the optimal e4CL-D5 condition, which generated 11.9% 8CLCs via inducing primed hESCs to a naïve state using 4CL conditions and then to 8CLCs using TSA and DZNep. Yoshihara (Yoshihara et al., 2022) reported the optimal condition for generating 8CLCs at 6.6% efficiency with a 15-minute *DUX4* pulse followed by a 12-hour incubation in a primed background, avoiding DUX4-mediated cytotoxicity (Bosnakovski et al., 2008). In contrast, TS used a 2–4-hour induction in their *DUX4*-inducible naïve human embryonic stem cells (hESC) for 8CLC generation. To ensure fair comparison, we exclusively utilized scRNA-seq datasets generated without prior enrichment for the 8CLC population. The TS DUX4 data was generated from nuclear RNA and thus we accounted for the inclusion of introns in the transcripts in the pre-processing. Additionally, sequencing data from the Yan et al. (Yan et al., 2013) study oocytes, zygotes, and embryos were obtained to provide a uniform embryo reference for comparison, along with (Bi et al., 2022) naïve and primed hESC samples to represent different hPSC backgrounds. We re-aligned all datasets to the GRCh38 human reference genome and quantified both gene and TE expression. We proceeded to downstream processing that included quality control, cell clustering and characterization of cell identities. After analyzing each dataset individually, we proceeded to integrating the datasets with the Yan embryo reference and readily processed embryo samples included Meistermann et al. study (Meistermann et al., 2021). Downstream analyses of the integrated dataset included clustering, interrogation of cluster identity based on gene and TE expression, and gene regulatory network (GRN) analysis.

**Figure 1.**
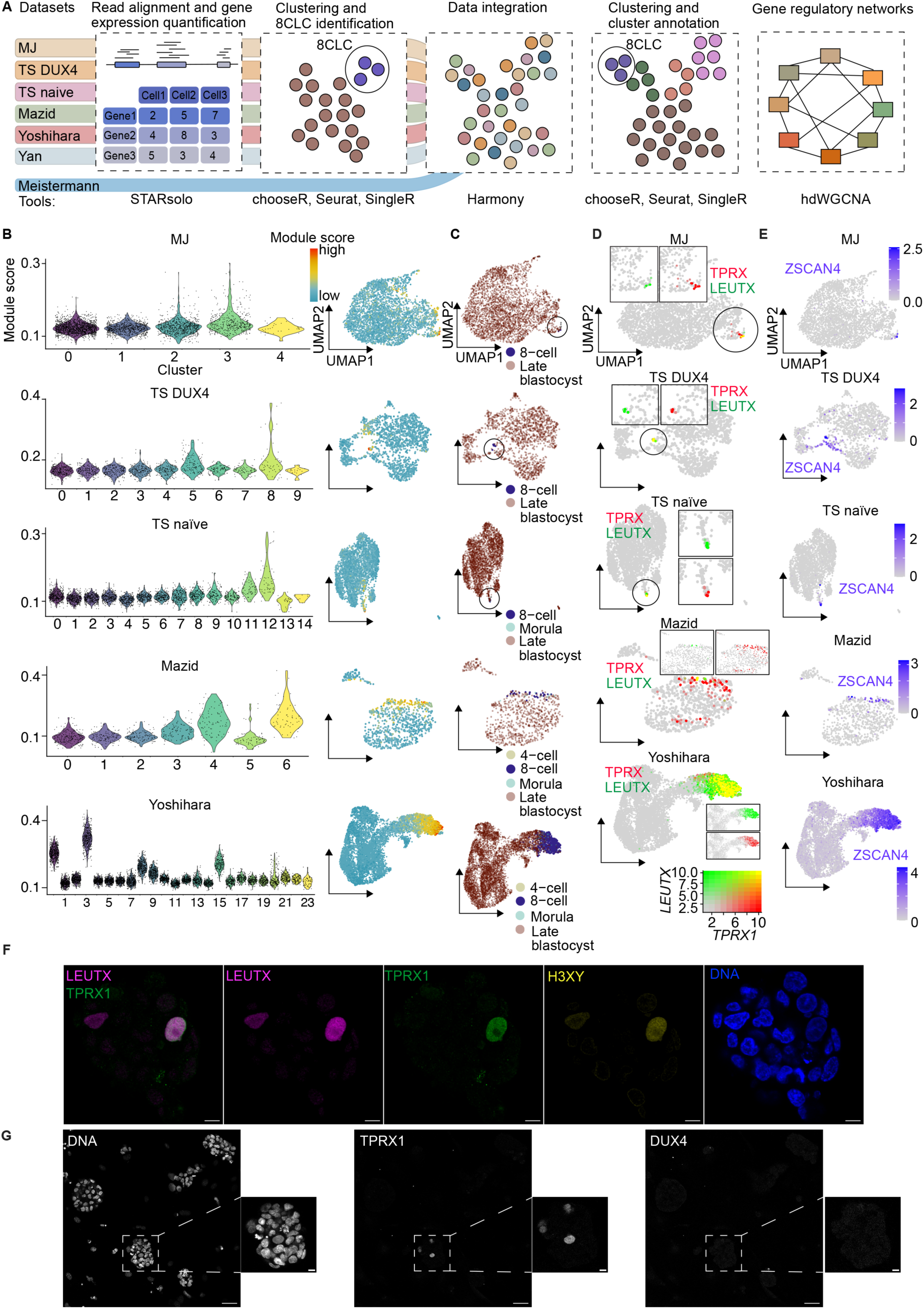
The expression of EGA genes distinguishes between intermediate and mature 8CLC identities. A) Schematic of the bioinformatics workflow and the used methods. B) Module scores for Stiparo et al. (2018) human 8-cell stage embryo-specific gene list across chooseR optimized (Fig. S1A) Seurat clusters (Fig. S1B) and as an UMAP of the hPSC datasets. Datasets: MJ, Moya-Jódar; TS DUX4, Taubenschmid-Stowers DUX4-induced; TS naïve, Taubenschmid-Stowers naïve. C) Results for Singler annotations with Yan et al. oocytes and embryos as reference projected as UMAPs of the hPSC datasets. From top to bottom: Moya-Jódar; Taubenschmid-Stowers DUX4-induced; Taubenschmid-Stowers naïve, Mazid, Yoshihara. D) Expression of *LEUTX* and *TPRX1* in the datasets. Colors represent single positive (red: *TPRX1*, green: *LEUTX*) and double positive (yellow) cells with 0.1 set as the threshold for expression. E) Expression of *ZSCAN4* in the datasets. The color intensity represents normalized and scaled expression. F) Naïve H9 hPSCs immunostained with antibodies recognizing LEUTX (magenta), TPRX1 (green), and H3XY (yellow). DNA counterstained with 4ʹ,6-diamidino-2-phenylindole. Scale bar 50 μm, in insets 10 μm. G) Naïve H9 hPSCs immunostained with antibodies recognizing TPRX1 (white) and DUX4 (white; not detected). DNA counterstained with 4ʹ,6-diamidino-2-phenylindole. Scale bars 10 μm.

To identify cells or entire clusters with resemblance to human 8-cell stage embryos, we clustered each dataset using the optimized resolution parameter determined by chooseR (Patterson-Cross et al., 2021) and visualized the results on a Uniform Manifold Approximation and Projection (UMAP) (Fig. S1A, Fig. S1B). We obtained lists of marker genes for human embryonic stages and lineages from an integrative transcriptome analysis by (Stirparo et al., 2018). Using these gene lists as reference for Seurat’s AddModuleScore function, we identified clusters with slightly elevated 8-cell stage embryo module scores in most datasets: clusters 11 and 12 for TS naïve, 5 and 8 for TS DUX4, 2 and 3 for MJ, and 4 and 6 for Mazid (Fig 1B). However, cells with the highest scores were outliers rather than representing the average cluster score, indicating that 8CLC identity was not the major driver of clustering. Contrarily, the Yoshihara dataset showed clear separation of clusters in terms of the 8-cell score, with clusters 0 and 3 having high scores and clusters 8, 9, and 15 having moderate scores. Additionally, in all datasets except TS DUX4, the early ICM scores had an opposite pattern to the 8-cell score, suggesting the loss of stem cell features in potential 8CLCs (Fig S1C).

Using the SingleR algorithm with re-processed Yan embryonic samples as reference, we annotated each cell to a developmental stage between oocyte and late blastocyst (Fig 1C). Notably, all datasets except Yoshihara had a maximum of 3.6 % of cells resembling 8-cell stage embryos, with over 800 cells, that is 11.4%, in Yoshihara (Table S2). Some cells in Mazid and Yoshihara datasets were also annotated as morulae, consistent with previous reports of some 8CLCs resembling morulae (Yoshihara & Kere, 2023; Zhao et al., 2024). Comparison of Stirparo 8-cell scores to SingleR annotations showed significantly higher scores in cells resembling 8-cell stage embryos compared to late blastocysts (FDR < 0.05, Wilcoxon rank sum test) (Fig. S1D). Interestingly, cells resembling 4-cell stage embryos and morulae had similar 8-cell scores as the 8-cell stage embryo annotated cells, indicating challenges in distinguishing 8CLCs from adjacent embryonic stages. Assessment of transitions in naïve versus primed cell state using Bi et al. (Bi et al., 2022) data as reference revealed that the majority of cells retained their original hPSC identity, except for TS DUX4, which showed a mixture of naïve and primed cell types, which may be due to DUX4 induction or technical factors (Fig. S1E). Additionally, cells annotated as 8-cell stage embryos in Yoshihara and Mazid datasets mainly retained their background hPSC identity with only some cells marked as unannotated in Yoshihara, indicating that the background hPSC identity remains identifiable in a majority of plausible 8CLCs. Since enrichment of cells in the G1 and G2M cell cycle phases have been reported in mouse 2CLCs (Atashpaz et al., 2020; Eckersley-Maslin et al., 2016; Zhu, Cheng, et al., 2021) we investigated whether this phenomenon could be observed in 8CLCs (Fig S1F). Analysis of cell cycle phase using Seurat’s CellCycleScoring function revealed that cells in G2M had significantly higher 8-cell scores compared to cells in S or G1 (FDR < 0.05, Wilcoxon rank sum test), suggesting conserved features regarding cell cycle regulation in mammalian EGA recapitulating cells.

An intermediate 2-cell-like state with heterogeneous 2-cell embryo stage marker gene expression has been reported in mice (Rodriguez-Terrones et al., 2018). We investigated the expression levels of 8-cell state marker genes, *TPRX1*, *LEUTX*, and *H3Y/X*, to determine if similar expression heterogeneity exists in humans. Notably, *TPRX1* is not a DUX4 target, while *H3Y/X* and *LEUTX* are. Moreover, *LEUTX* is suggested to be a specific marker for human 8-cell stage embryos (Yoshihara et al., 2022). We confirmed that this gene combination is a minimum requirement for identifying 8CLCs, as all Yan 8-cell stage embryos expressed these genes (Table S3). However, most Yan morulae were also triple-positive, indicating these markers are not exclusive to the 8-cell stage (Yan et al., 2013). In the 8CLC datasets, 0.17% to 49.6% of cells were triple-positive, with the lowest and highest fractions in the MJ and Yoshihara datasets, respectively (Table S4). Most datasets had similar percentages of double- and triple-positive cells, but in the Mazid dataset, *LEUTX* upregulation limited 8CLC generation: *TPRX1* and *H3Y/X* double-positive cells accounted for 11.3% of all cells, compared to 1.15% of triple-positive cells. In contrast, the Yoshihara dataset showed no such limitation, with four times as many triple-positive cells as SingleR-annotated 8CLCs. *TPRX1* and *LEUTX* double-positive cells overlapped with the highest Stirparo 8-cell scores and matched SingleR 8-cell stage embryo annotations in all datasets (Fig. 1D). These cells also showed the highest *ZSCAN4* expression (Fig. 1E), while *DUX4* expression was limited to a few cells in most datasets, except Yoshihara, where it overlapped with 8-cell embryo annotations (Fig. S1G). We confirmed the presence of triple-positive cells at protein level by immunostaining naïve H9 hESCs with antibodies against TPRX1, H3X/Y, and LEUTX (Fig. 1F), which are also detected in human 8-cell stage embryos (Jouhilahti et al., 2016; Taubenschmid-Stowers et al., 2022). Notably, DUX4 protein was undetectable in naïve H9 hESC cultures (Fig. 1G). Approximately 5% of mESCs are ZSCAN4-positive, but only about 0.1-0.4% belong to the 2CLC population (Falco et al., 2007; Genet & Torres-Padilla, 2020). Single *ZSCAN4*-positive cells lacking *MERVL* expression exhibit a transcriptional profile between that of mESCs and *ZSCAN4/MERVL* double-positive 2CLCs (Rodriguez-Terrones et al., 2018). Similarly, MJ observed that only 0.3% of all naïve cells were double-positive for *ZSCAN4* and *MLT2A1*, indicating a true 8-cell-like identity. TS reported that while *ZSCAN4* is upregulated in most 8CLCs, markers like *TPRX1* or *DUXA* show less widespread expression. This, along with our findings, suggests an intermediate 8CLC state likely exists in humans, alike to mouse 2CLCs.

Based on the comparison of Stirparo 8-cell scores, marker gene expression, and SingleR annotations, we considered cells annotated as 8-cell stage embryos by SingleR as 8CLCs, with the Yoshihara and Mazid datasets having 11.4 and 3.6% of 8CLCs, respectively, and thus performing best. Most datasets showed fewer 8CLCs than originally reported, likely due to others using non-discriminating markers or different clustering approaches to annotate 8CLCs. Our results may contradict the original numbers partly because we introduced the concept of intermediate 8CLCs — 8-cell-like cells expressing only a subset of central marker genes. Discrepancies exist regarding the presence of an 8CLC subpopulation in 5iLA and t2iLGo naïve hPSCs. While TS and Mazid reported such populations, Yoshihara and Kere (Yoshihara & Kere, 2023) and Yu et al (X. Yu et al., 2022) did not confirm this, noting only low expression levels of some 8C genes in rare cells. These variations may stem from different scRNA-seq datasets and criteria used to distinguish 8CLCs. Based on our own and previous findings, we propose that stringent annotation criteria are essential when determining the existence of a mature 8CLC population sufficient to serve as an *in vitro* model of human EGA.

### Integrated analysis of the hPSC datasets and human embryos

To further explore our initial findings, we used Harmony (Korsunsky et al., 2019) to integrate the 8CLC datasets with the re-processed Yan et al. (2013) and the processed Meistermann et al. (2021), Fogarty et al. (2017), and Petropoulous et al. (2016) human oocyte and embryo data. We excluded the Fogarty et al. *OCT4* knockout embryos due to their reported developmental phenotype. Visual inspection of the batch-corrected integration result projected on a UMAP confirmed sufficient developmental stage-wise overlap for the Meistermann and Yan embryo samples and the two biological replicates of the Yoshihara and Mazid datasets (Fig. 2A, Fig. 2B). The embryonic samples were segregated from a majority of the hPSCs, which mainly grouped according to the naïve or primed hPSC background (Fig. 2C). Seurat clustering with 20 principal components (PCs) and the chooseR-optimized resolution parameter of 2.0 generated 29 clusters (Fig. 2D). We noted minor inter-cluster differences in the fractions of mitochondrial transcripts, with embryo-enriched clusters showing slightly elevated levels, likely due to less strict filtering of these samples (Fig. S2A). Conversely, the fraction of ribosomal protein-encoding transcripts showed high intra- and inter-cluster variation, which may be due to cell viability or biological differences (Fulka et al., 2020; Fig. S2B).

**Figure 2.**
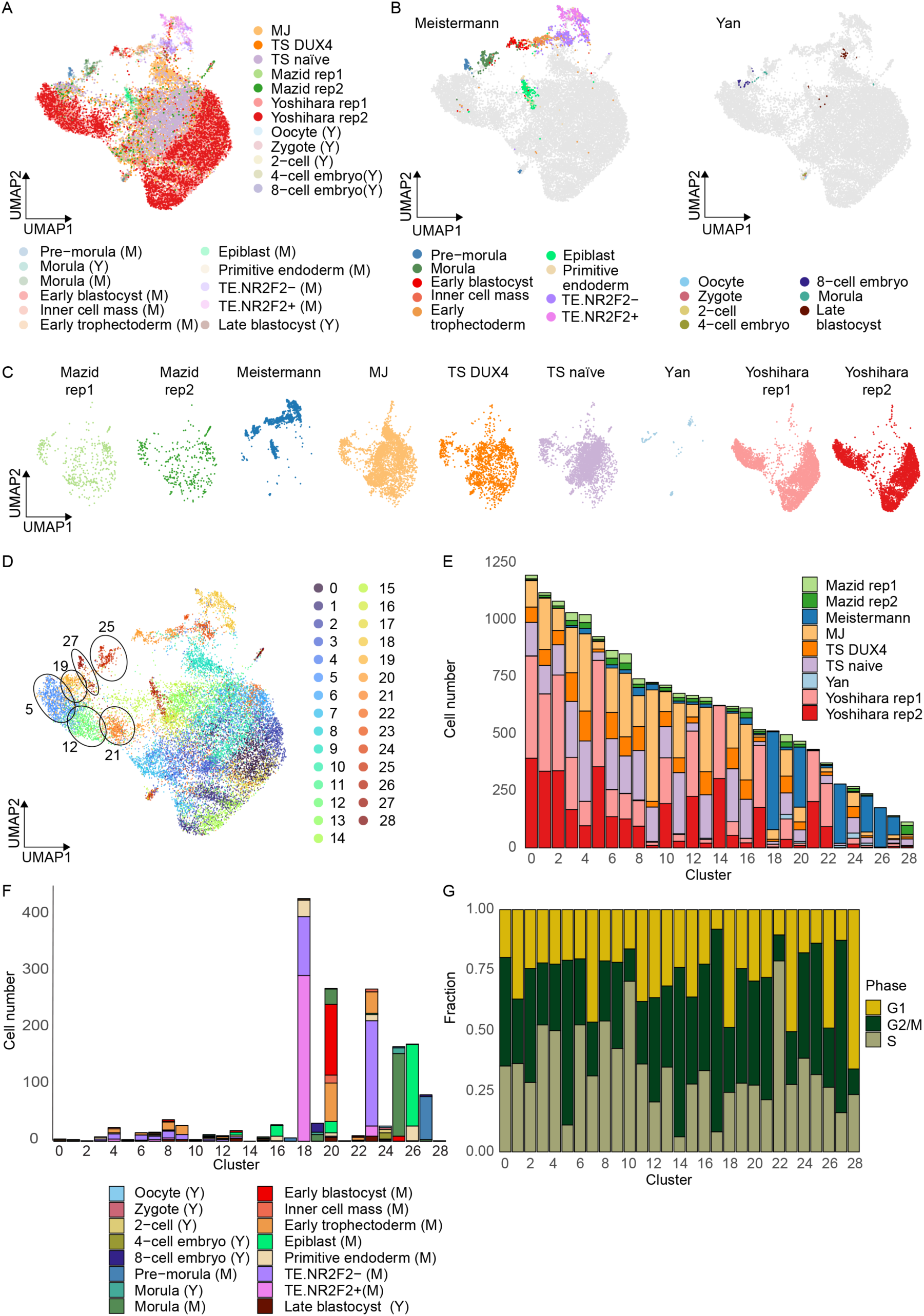
Integrated analysis of the hPSC datasets and human embryos. A) UMAP of the integrated dataset consisting of the hPSC datasets as well as human oocyte and embryo datasets, B) Meistermann et al. (2021) (left) and Yan et al. (2013) (right), C) and split by each original dataset and replicate. Datasets: MJ, Moya-Jódar; TS DUX4, Taubenschmid-Stowers DUX4-induced; TS naïve, Taubenschmid-Stowers naïve; M, Meistermann; Y, Yan. Abbreviations: rep, replicate. D) Seurat clusters of the integrated dataset using 20 PCs and resolution 2.0. For resolutions 1.8 and 2.2 see Fig. S2C. E) The number of cells from each original dataset and replicate and F) from the different embryonic stages in the integrated clusters. G) The fraction of cells in each cell cycle phase in the integrated clusters. See Fig. S2D for information on enriched phase for each cluster.

We calculated the numbers of hPSCs from each 8CLC dataset (Fig. 2E, Table S5) as well as the number of oocytes and embryos in each cluster. Most Yan 8-cell stage embryos and Meistermann pre-morulae, the corresponding stage, were included in clusters 19 and 27, respectively (Fig. 2F, Table S5). hPSCs from all datasets formed the major fraction of cells in cluster 19, while cluster 27 consisted mainly of Meistermann pre-morulae. We compared resolutions 1.8 and 2.2 with 2.0 to study their effects on confinements of clusters containing 8-cell stage embryos (Fig. S2C). At resolution 1.8, hPSCs included previously in cluster 27 with the Meistermann pre-morulae were relocated to a neighboring cluster that now consisted of Yan 8-cell stage embryos and hPSCs. At resolution 2.2, cluster 27 expanded to include hPSCs from the neighboring cluster. Thus, the cluster boundaries showed small alterations upon changes in the resolution parameter in the area with the 8-cell stage embryos, yet the region as an entity remained rather stable. Finally, we inspected the cell cycle phase of all cells and noted variation in the predominant phase between the clusters (Fig. 2G). Clusters 10 and 22, which consisted of hPSCs and trophectoderm cells, were enriched for the S phase, while none of the clusters showed substantial enrichment for the G1 phase (p-value < 0.05, Chi-square test of independence, Fig. S2D). Interestingly, the G2M phase was enriched in clusters 5, 14, and 17, which consist primarily of Yoshihara hPSCs. Cluster 5 is proximal to the 8-cell stage embryos, suggesting that *DUX4* induction may promote cell cycle changes coupled with the emergence of 8CLCs.

### Intermediate 8CLCs retain stem-cell like characteristics absent in human 8-cell stage embryos

To annotate the clusters, we calculated module scores using gene expression signatures specific to naïve and primed hPSCs as well as embryonic stages. We used gene lists from studies on the transition from naïve to primed hESCs (Bi et al., 2022) and consensus markers across multiple hPSC lines (Messmer et al., 2019), as references for hPSC identities. The naïve and primed modules showed opposite patterns aligned with hPSCs background, as expected (Fig. 3A). While Yoshihara hPSCs generally showed high primed scores, clusters 5 and 12 that contain mainly Yoshihara hPSCs, had lower primed scores (Fig. 3B). Since naïve cells are reported to spontaneously generate 8CLCs, we hypothesized that a naïve background might be more conducive to 8CLC identity. Generally, embryonic samples exhibited a naïve-like profile (Fig. 3C, Fig. S3A), matching findings by Yoshihara et al. (2023) and aligning with reports of spontaneous EGA gene upregulation in naïve but not primed hPSCs.

**Figure 3.**
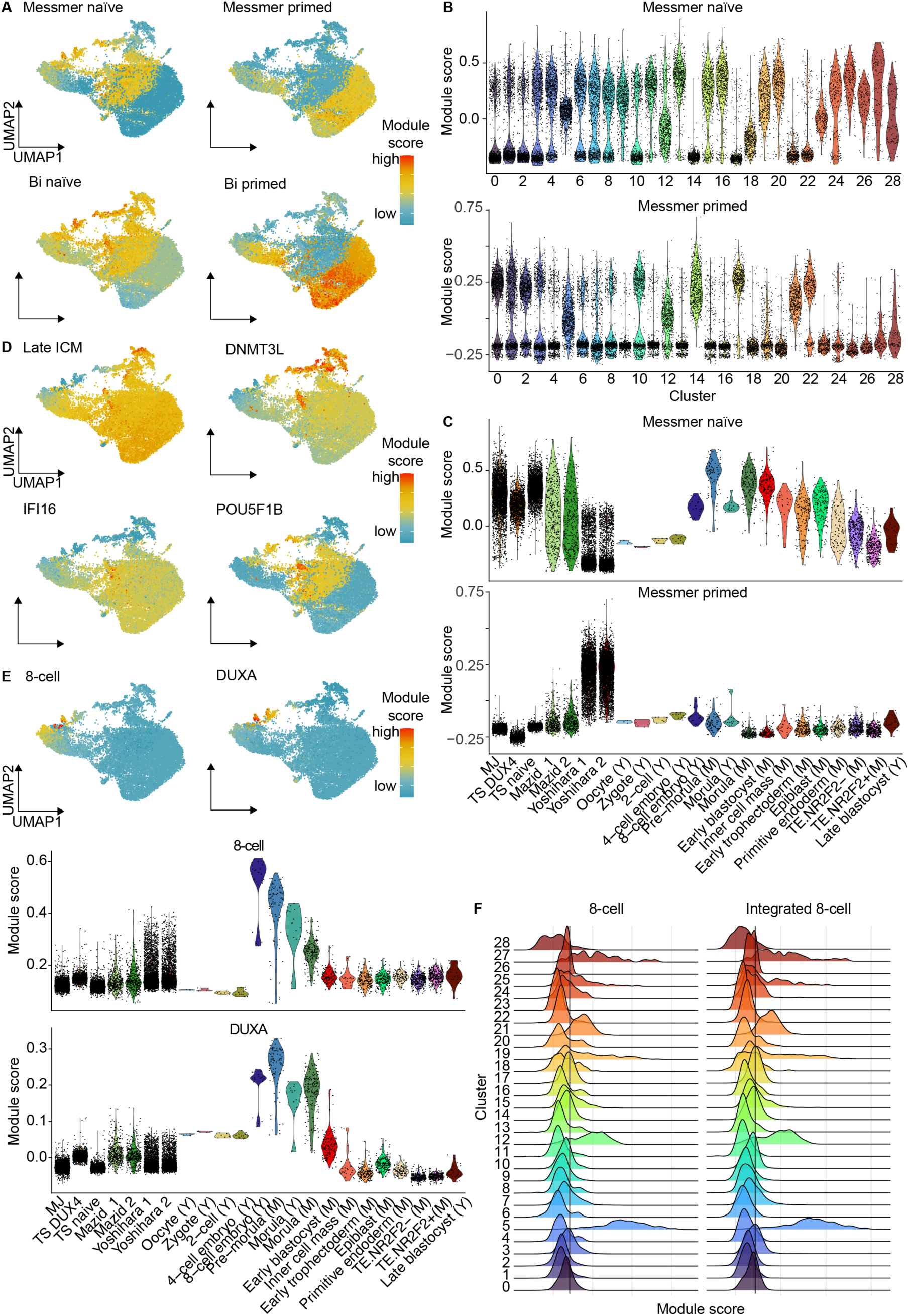
Intermediate 8CLCs retain stem-cell like characteristics absent in human 8-cell stage embryos. A) Module scores of Messmer et al. (2019) (above) and Bi et al. (2022) (below) naïve (left) and primed (right) hPSC gene lists as an UMAP of the integrated dataset. B) Module scores of Messmer et al. (2019) naïve and primed hPSC gene lists in the integrated dataset, across clusters and C) across the original datasets and replicates, as well as the embryonic stages in Meistermann et al. (2021) and Yan et al. (2013). Datasets: MJ, Moya-Jódar; TS DUX4, Taubenschmid-Stowers DUX4-induced; TS naïve, Taubenschmid-Stowers naïve; M, Meistermann; Y, Yan. D) Module scores of Stirparo et al. (2018) (Late ICM) and Meistermann et al. (2021) (DNMT3L, IFI16, POU5F1B) late embryonic stage gene lists and E) Stirpro et al. (2018) (8-cell) and Meistermann et al. (2021) (DUXA) 8-cell stage embryo specific gene lists in the integrated dataset as an UMAP (top), and across the original datasets and replicates, as well as the embryonic stages in Meistermann et al. (2021) and Yan et al. (2013) (bottom). For other embryonic modules see Fig. S3B. F) Distributions of module scores calculated prior to (left) and after (right) integration for Stirparo et al. (2018) 8-cell stage gene list across the integrated clusters. The vertical line depicts the mean value.

We evaluated the identity of the clusters using embryonic stage and lineage-specific gene expression signatures from Meistermann et al. (2021) and Stirparo et al. (2018). Several modules were highly specific to early embryonic stages other than the 8-cell stage (Fig. S3B). Early-stage modules, including Stirparo zygote and 4-cell, and Meistermann ZSCAN4, scored highest in oocytes to 4-cell embryos (cluster 24) and moderately in 8-cell embryos (cluster 27). Modules related to morula-early blastocyst stages, Stirparo morula and early ICM, were elevated in hPSCs but peaked in morulae and post-morula stages or lineages. Similarly, lineage-specific modules, Meistermann GATA4, GATA2, and NR2F2, had high scores in primitive endoderm (PrE) and trophectoderm cells, as expected.

We identified several modules indicative of stem cell-like gene expression, referred to as blastocyst-hPSC modules, with high scores in stages succeeding 8-cell embryos and in hPSCs (Fig. 3D). The Stirparo late ICM and DNMT3L modules had high scores in post-morula stage cells and hPSCs, except for some cluster 5 hPSCs (Fig.3D). Additionally, the IFI16 module scored high in most hPSCs, epiblast, PrE, and late blastocyst cells but low in other embryonic samples and some cluster 5 hPSCs, indicating that these hPSCs lack typical stem cell-like gene expression (Fig.3D). The POU5F1B module scored highest in morulae, early blastocyst, epiblast, and late blastocyst cells, as well as naïve hPSCs, while oocytes to 8-cell embryos and Yoshihara hPSCs scored lowest (Fig.3D). These findings suggest that despite clustering with 8-cell stage embryos, many hPSCs in clusters 19 and 27 retain stem cell-like characteristics, while hPSCs in cluster 5 have lost them. Finally, the 8-cell stage embryo modules, Stirparo 8-cell stage and Meistermann DUXA, scored highest in Yan and Meistermann 8-cell embryos and morulae and moderately in several hPSCs in clusters 5, 19, and 27, indicating these clusters may harbor hPSCs with varying levels of 8CLC identity (Fig. 3E).

Finally, we compared initial dataset-wise and re-calculated integration-based Stirparo 8-cell module scores, revealing slightly elevated scores for clusters 12, 19, and 21 before and after integration (Fig. 3F). Clusters 5 and 27 had clearly above-average scores. Most hPSCs identified as 8CLCs in dataset-wise analyses were in cluster 5 of the integrated data (Fig. S3C), rather than cluster 27. Consequently, we conclude that while most 8-cell embryos reside in clusters 19 and 27, the majority of 8CLCs reside in cluster 5, with possibly some in cluster 27. Clusters 12, 21, and 19 likely contain intermediate 8CLCs, showing some 8-cell stage-specific gene upregulation but insufficient downregulation of stem cell-like gene expression, similar to previously reported DUX4-induced intermediate 8CLC and mouse intermediate 2CLCs (Rodriguez-Terrones et al., 2018; Yoshihara et al., 2022).

### TE expression can supplement gene expression data to distinguish between intermediate and mature 8CLCs

The expression of non-coding loci, including repeat elements, during early development is conserved across model organisms and humans. TEs function as promoters and enhancers and produce RNAs and proteins that are likely functional (Y. Guo et al., 2024; Peaston et al., 2004; Vuoristo et al., 2022; Whiddon et al., 2017). In mice, MERVL expression distinguishes intermediate from fully mature 2CLCs (Rodriguez-Terrones et al., 2018), highlighting the importance of TEs in this context. Despite observations of TE expression in 8CLCs, a formal comparison of TE expression in different 8CLC populations is lacking. We used the scTE algorithm to quantify TE expression in 8CLC datasets and Yan oocyte and embryo samples. We used Liu et al. (2017) human oocyte to blastocyst TE expression data as reference and identified 12 TE expression trajectories during early human development using fuzzy clustering (Fig. 4A). We curated these expression trajectories to delineate 8-cell embryo-like TE expression signatures (Fig. 4B, Fig. S4A) and calculated module scores for our integrated dataset, excluding Meistermann samples due to lack of scTE processing. Most signatures were recapitulated in Yan oocytes and embryos, except the 8-cell negative signature (fuzzy cluster 6) (Fig. 4C, Fig. S4B).

**Figure 4.**
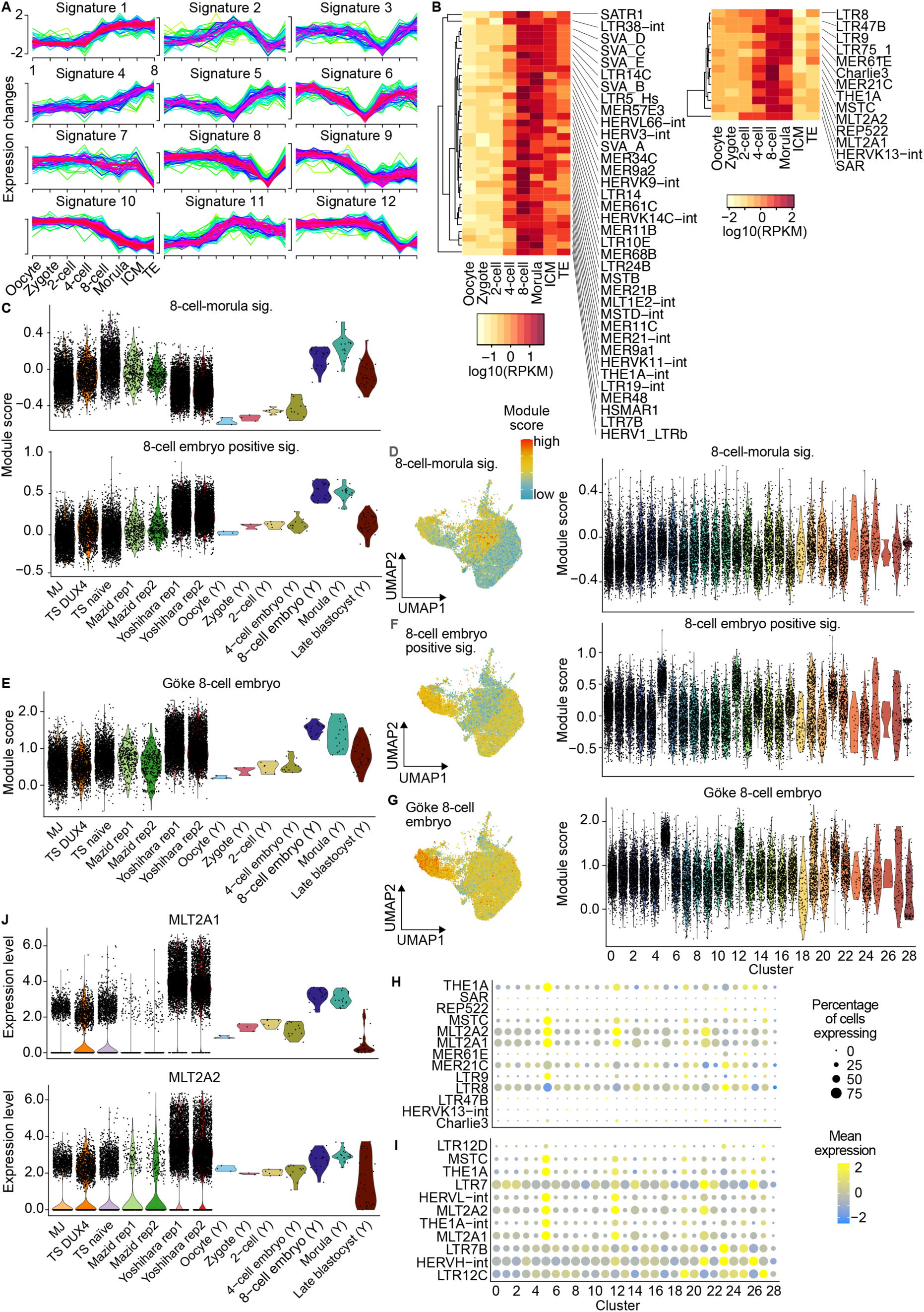
TE expression can supplement gene expression data to distinguish between intermediate and mature 8CLCs. A) Fuzzy clustering of TE expression in human oocytes and early embryos in Liu et al. (2019). B) Hierarchically clustered (Euclidean distance and average linkage) heatmaps of TE expression in human oocytes and early embryos (Liu et al. 2019) for the 8-cell-morula (fuzzy cluster 11; left) and the 8-cell positive signatures (fuzzy cluster 2; right). C) Module scores of the 8-cell-morula and 8-cell positive signature in the hPSC datasets and replicates, as well as the embryonic stages in Yan et al. (2013). Datasets: MJ, Moya-Jódar; TS DUX4, Taubenschmid-Stowers DUX4-induced; TS naïve, Taubenschmid-Stowers naïve; Y, Yan. D) Module scores of the 8-cell-morula signature as a UMAP (left) of the integrated dataset and across the clusters (right). E) Module scores of Göke 8-cell stage embryo TE list in the hPSC datasets and replicates, as well as the embryonic stages in Yan et al. (2013). F) Module scores of the 8-cell positive signature, G) and Göke 8-cell stage embryo signature, as an UMAP (left) of the integrated dataset and across the clusters (right). H) Dotplot of TE expression across clusters in the integrated dataset with cluster average expression (scaled) indicated in color and percentage of cells expressing TE as dot size for the 8-cell positive and I) Göke 8-cell stage embryo signatures. J) Expression of MLT2A1/2 in the hPSC datasets and replicates, as well as the embryonic stages in Yan et al. (2013).

The morula-blastocyst signature (fuzzy cluster 1) scored highest in late blastocysts and naïve stem cell clusters, especially cluster 19 with naïve intermediate 8CLCs (Fig. S4C), suggesting that these cells exhibit TE expression typical of 8-cell embryo succeeding stages. Primed-background intermediate 8CLCs and 8CLCs in clusters 5, 12, and 21 showed lower scores for this signature. The scores were driven by Alu-element expression, which have also been linked to human EGA (Töhönen et al., 2015), while most cells lacked LTR expression (Fig. S4D). The 8-cell-morula signature (fuzzy cluster 11) scored highest in 8-cell embryos, morulae, naïve hPSC clusters, and intermediate 8CLCs and 8CLCs in clusters 19 and 5 (Fig. 4D), though only a fraction of TEs in this signature were expressed (Fig. S4E) suggesting that it may be specific to the Liu et al. dataset due to biological or technical reasons.

We used the 8-cell positive (fuzzy cluster 2) and an 8-cell specific TE signature from Göke et al. (2015) to validate the identity of putative 8CLCs in terms of TE expression (Fig. 4E). Out of the studied embryonic stages both signatures showed the highest scores in 8-cell stage embryos and morulae (Fig. 4F-G). Clusters 5, 12, and 21, mainly containing Yoshihara 8CLCs and intermediate 8CLCs, scored highest, with some cells in clusters 19 and 27 also showing high scores (Fig. 4H-I), suggesting they may also possess 8-cell-like identity in terms of TE expression. MLT2A1 and MLT2A2, included in both signatures, were highly expressed in cluster 5 8CLCs, as expected, being 8-cell specific and DUX4 targets (Geng et al., 2012; Hendrickson et al., 2017; Liu et al., 2019; Young et al., 2013) (Fig. 4J). While some TEs in the 8-cell positive signature were absent in most cells, 8CLCs in cluster 5 expressed all Göke 8-cell specific TEs except LTR12D. Overall, these results confirm cluster 5 cells and some cells in cluster 27 are likely 8CLCs, while hPSCs in clusters 12, 19, and 21 are intermediate with incomplete 8CLC gene and TE expression. Moreover, TE expression in cluster 19 intermediate 8CLCs resembles that of later embryonic stages, suggesting that TE expression could be used to distinguish intermediate 8CLCs from mature ones in humans, similar to mouse 2CLCs. It is tempting to speculate that induced upregulation of early factors like *DUX4* in hPSCs (Vuoristo et al., 2022) may be crucial for rewiring non-coding elements involved in EGA networks. Other PRD-like homeobox transcription factors such as LEUTX (Gawriyski et al., 2023) and OBOX (Ji et al., 2023), are also reported to regulate TEs as activators, while DUXA (Bosnakovski et al., 2023) and OTX2 (Guo et al., 2021; Liu et al., 2019) act as repressors in mammals. Activation of the non-coding genome, including TEs, is likely crucial for 8CLC generation, possibly mediated by demethylases, methylases, and acetylase inhibitors like DPPA3 (Huang et al., 2017) and KDM4 which have been implicated in chemical 8CLC generation (Mazid et al., 2022; X. Yu et al., 2022). However, active TEs may cause cell toxicity in long-term culture, complicating sustained cell viability.

### GRN analysis indicates that changes in energy and RNA metabolism, alternative splicing, and ribosome biogenesis accompany the emergence of 8CLCs

We conducted GRN analysis using hdWGCNA (Morabito et al., 2023), a single-cell adapted form of weighted gene co-expression network analysis (WGCNA, (Langfelder & Horvath, 2008)), to identify the GRNs governing the 8CLC state. We focused on differences between intermediate 8CLCs to highlight reprogramming barriers to achieving the 8-cell-like state. We grouped Seurat clusters based on inferred annotations (Fig. 5A, Fig. S5A-B). Clusters containing specific embryonic stages or lineages were named according to their embryonic identity in the Yan or Meistermann datasets (Fig. S5C). Cluster 5, primarily composed of Yoshihara 8CLCs, was named Yoshihara-8CLC, while clusters 12 and 21, as well as 19, containing intermediate 8CLCs, were named Yoshihara-intermediate and 8-cell-like, respectively. Non-8CLC hPSC clusters were categorized as Naive-like, Primed-like, and Mixed-like and cluster 28 was defined as an outlier due to its minimal resemblance to other clusters.

**Figure 5.**
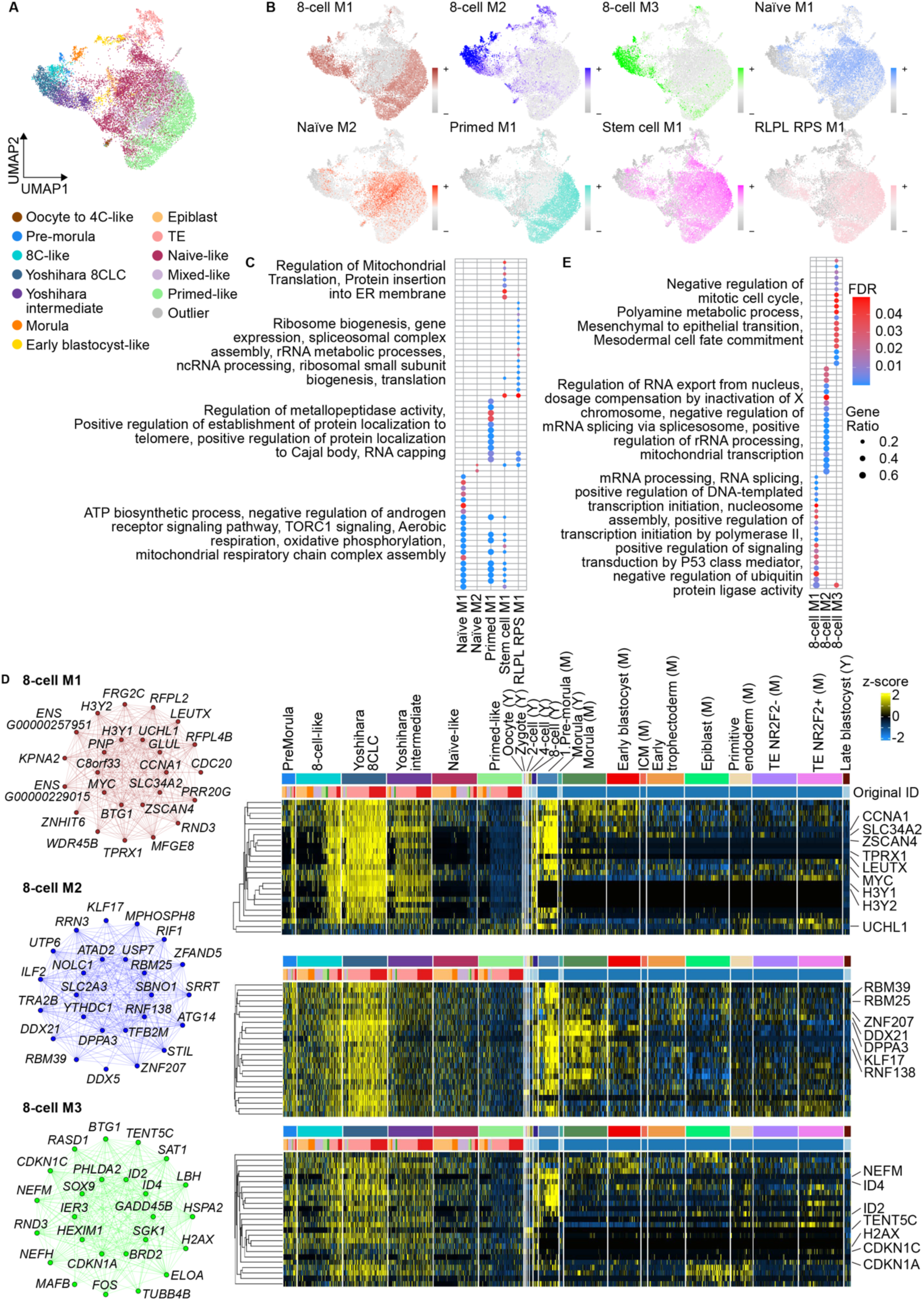
GRN analysis indicates that changes in energy and RNA metabolism, alternative splicing, and ribosome biogenesis accompany the emergence of 8CLCs. A) UMAP of the integrated data with cell groups used for GRN analysis indicated in colors. Clusters have been combined to bigger entities based on similarity as depicted in Fig. S5B. B) Curated hdWGCNA gene co-expression networks on UMAPs of the integrated dataset with the harmonized module eigengene scores indicated by the color intensity. C) Enriched biological process gene ontology terms for modules representative of stem cell-like states. Top 20 terms by gene ratio and FDR < 0.05 are shown for each GRN. D) Network plots (left) and hierarchically clustered (Euclidean distance and average linkage) expression heatmaps (right) of hub genes GRNs representative of intermediate and mature 8CLCLs and 8-cell stage embryos. Heatmap colors indicate z-scores that have been calculated in a dataset- and replicate-wise manner. See Fig. S5D for corresponding results for stem cell GRNs. Abbreviations: ICM, inner cell mass; TE, trophectoderm; M, Meistermann et al.; Y, Yan et al. E) Enriched biological process gene ontology terms for modules representative of intermediate and fully mature 8CLCLs and 8-cell stage embryos. Top 20 terms by gene ratio and FDR < 0.05 are shown for each GRN.

We identified eight co-expression modules informative in distinguishing 8-cell-like states from non-8CLC hPSCs using manual curation. Four modules related to general stem cell identity and naïve or primed cell backgrounds (Fig. S5D). The Stem cell-M1 module had high scores in non-8CLC hPSCs and intermediate 8CLCs, featuring hub genes coding for mitochondrial complex (NDUF) and ribosomal proteins, which decreased along the trajectory from hPSCs to intermediate 8CLCs to 8CLCs. The Naïve-M1 and Naïve-M2 modules scored highest in intermediate 8CLCs and naïve hPSCs, featuring the hub gene UPP1, which a previous study reported as lowly expressed in Yoshihara DUX4-induced 8CLCs compared to other 8CLCs (Yoshihara & Kere, 2023). We find that the other 8CLCs also downregulate this gene while intermediate 8CLCs, from both Yoshihara and other datasets do not, highlighting that the original annotation of the cells as intermediate or mature 8CLCs affect downstream comparisons. Contrarily, IFITM1, enriched in epiblast, PrE, hPSCs, intermediate 8CLCs, and a subset of naïve 8CLCs but not Yoshihara 8CLCs, is upregulated on the transcript and surface protein level in human naive hPSCs compared to primed hPSCs (Grow et al., 2015). It may contribute to the immunoprotective role of HERVK against viral infections in human embryos and germ cells. Finally, the Primed-M1 module was highly expressed in some intermediate 8CLCs and primed non-8CLC hPSCs, as expected.

GO analysis of biological processes associated with these modules highlighted the formation of the mitochondrial ATP pathway, indicating that energy metabolism distinguishes stem cells and intermediate 8CLCs from mature 8CLCs and 8-cell stage embryos (Fig. 5C). Previous studies suggest that 8CLC generation might be modulated through metabolic interference similar to mouse 2CLCs, which exhibit lower mitochondrial respiratory capacity, reduced oxygen consumption rates, and reduced glycolytic activity compared to mESCs (Genet & Torres-Padilla, 2020; Hu et al., 2020; Mazid et al., 2022). Moreover, mammalian early embryos primarily rely on oxidative phosphorylation for energy metabolism and display lower oxygen consumption compared with blastocysts (Dumollard et al., 2009; Gardner & Wale, 2013; Genet & Torres-Padilla, 2020). Our integrative GRN analysis suggests that downregulation of mitochondrial respiration occurs during 8CLC generation, warranting further investigation to whether it can be modulated to generate 8CLCs, and if it involves remodeling of mitochondrial structures as observed in mouse 2CLCs (Genet & Torres-Padilla, 2020).

We discovered a ribosomal protein (RP) gene module, RLPL-RPS-M1 with high scores in a subset of intermediate 8CLCs and many hPSCs, (Fig S5D). This module’s hub genes, which included many RP coding genes, were expressed at low levels in embryos until the morula stage, and 8CLCs (clusters 27 and 5), indicating downregulation of RP gene expression in 8CLCs and 8-cell stage embryos compared to stem cells and later preimplantation embryonic stages. Previous studies have linked the disruption of rRNA synthesis with the induction of the 2C state (Chen et al., 2021; Percharde et al., 2018; Xie et al., 2022; Yu et al., 2021). Recent findings reveal that RPs are generally expressed at low levels right before the transition from mESCs to 2CLCs, indicating their potential role as barriers in this transition (Yi et al., 2023; Zhu, Cheng, et al., 2021). Knockdown of certain RPs in mESCs induced transcriptome and chromatin accessibility profiles resembling 2CLCs, with Rpl14 depletion leading to decreased TRIM28 binding to Dux and rRNA loci, suggesting it may act upstream of Dux. Additionally, we found that DUX4 induction resulted in changes in RP expression, indicating potential bidirectional crosstalk. GO analysis associated terms related to ribosome biogenesis and translation with this GRN module, as expected. These findings collectively suggest that different types of 8CLCs and early embryos are in a quiescent state compared to metabolically active stem cells and later embryonic stages, providing insights into generating and possibly maintaining the 8CLC state.

The 8-cell-like state was demarcated by three gene co-expression modules (Fig 5D). The 8-cell-M2 module exhibited high scores in 8-cell stage embryos, morulae, and a diverse array of 8-cell stage embryo-resembling cells, encompassing both intermediate and mature 8CLCs from both naïve and primed backgrounds. Hub genes within this module included markers like *DPPA3* and *KLF17*, previously associated with the spontaneous and chemically induced emergence of 8CLCs in hPSCs, suggesting their essential role in facilitating 8CLC emergence without genetic induction of minor EGA genes. Studies have consistently reported upregulation of *KLF17* and *DPPA3* in 8CLCs compared to hESCs, with *KLF17* levels peaking at the 8-cell stage in human embryos, underscoring its potential as a key player in 8CLC generation (Blakeley et al., 2015; Taubenschmid-Stowers & Reik, 2023; Yoshihara & Kere, 2023). Furthermore, enrichment of KLF17 binding motifs around 8CLC genes and its involvement in 8CLC GRNs alongside DPPA3 suggest their central roles in 8CLC biology, supported by findings of DPPA3 knockout downregulating EGA gene expression (Mazid et al., 2022). The expression levels of hub genes formed a strengthening gradient along the axis from non-8-cell stage embryo samples to intermediate 8CLCs to mature 8CLCs and 8-cell stage embryos. Interestingly, both intermediate 8CLC clusters 12 and 19, predominantly comprising primed and naïve hPSCs, respectively, exhibited intermediate and heterogenous expression levels of the hub genes. This suggests that naïve cells, Mazid medium-induced naïve-like cells, and DUX4-induced naïve and primed cells could utilize similar pathways to reach the 8CLC state.

GO analysis revealed that the 8-cell-M2 module was enriched for terms related to RNA metabolism and splicing. Recent studies have highlighted disruptions in splicing during mammalian and human EGA, resulting in novel non-coding isoforms and inclusion of ORF-disrupting exons in transcripts (Torre et al., 2023). Moreover, splicing inhibition can induce an EGA-like state in both mESCs and hESCs (Li et al., 2024; Shen et al., 2021; Taubenschmid-Stowers & Reik, 2023) and Mazid et al. (2022) observed that many 8CLC genes are associated with GO categories related to protein synthesis and RNA metabolism, consistent with the enrichment of these processes in the 8-cell stage embryo (Stirparo et al., 2018). Our findings, along with other studies, suggest that these processes are central to generating 8CLCs, both spontaneously and via genetic methods, as well as to normal embryonic development.

The 8-cell-M1 module exhibited the highest scores in DUX4-treated primed hPSCs, intermediate and mature 8CLCs, as well as 8-cell stage embryos. Hub genes in this module included known 8-cell stage embryo markers, such as *TPRX1*, *H3Y1*, *H3Y2*, *ZSCAN4*, *CCNA1*, LEUTX, and *SLC34A2*, many of which are downstream targets of DUX4 (Geng et al., 2012; Hendrickson et al., 2017; Resnick et al., 2019; Vuoristo et al., 2022). While the expression patterns of many genes were similar between 8-cell stage embryos and cluster 5 8CLCs, confirming DUX4’s potency in upregulating EGA expression (Nykänen & Vuoristo, 2024), the module may lack EGA genes that are not downstream of DUX4. Higher scores of the Stirparo 8-cell stage module in embryos compared to 8CLCs suggest that induction factors beyond DUX4 are needed for accurate i*n vitro* models of 8-cell stage embryos. Interestingly, *UCHL1*, expressed in Yan minor EGA stage and Meistermann 8-cell stage embryos, was more prevalent in Yoshihara intermediate 8CLCs than mature 8CLCs, suggesting its role in 8CLC generation. Embryos from *UCHL1* mutant female mice fail to undergo morula compaction and do not form blastocysts *in vivo,* indicating a maternal effect related to *UCHL1* deficiency (Mtango et al., 2012). Recently, Zou et al. (2022) found that PRD-like homeobox factors *TPRX1*/*2*/*L* became highly translated before or during EGA, and their triple knockdown downregulated several EGA transcripts, impairing human preimplantation development. Furthermore, Mazid et al. (2022) showed that knockout of *TPRX1* impaired 8CLC generation, indicating parallel routes such as *DUX4* or *TPRX1*/*2*/*L* induction exist upstream of not only 8-cell stage embryos but also their *in vitro* models.

The 8-cell-M1 module is enriched in GO terms associated with p53-mediated signaling, negative regulation of ubiquitin protein ligase activity, and stem cell proliferation (Fig. 5E). The p53 pathway, downstream of hub gene MYC the mRNA of which is stabilized upon *DUX4* expression (Shadle et al., 2017), is activated by DNA damage and replication stress, linked to the transition from mESCs to 2CLCs (Atashpaz et al., 2020; Grow et al., 2021; Nakatani et al., 2022). Yu et al. showed that the p53 activator CBL0137 enhances the expression of 8CLC genes in hPSCs. Recent findings suggest *DUX4* activation by p53 (Grow et al., 2021), potentially indicating an autoregulatory loop involving DUX4, MYC, and p53.

While p53 activation can induce the 2C-like state upon DNA damage, it may not be necessary for Dux de-repression in response to nucleolar disruption (Bowling et al., 2018; Percharde et al., 2018), highlighting potential parallel pathways governing Dux/DUX4 expression in mammalian EGA. Additionally, the MYC-mediated apoptotic pathway and the double-stranded RNA innate immune response pathway serve as mediators of DUX4-induced apoptosis (Shadle et al., 2017), suggesting a role for MYC in the limited long-term culture support for DUX4-induced 8CLCs and possibly also other 8CLCs.

The 8-cell-M3 module exhibited the highest scores in cluster 5 8CLCs and 8-cell stage embryos. Several hub genes were expressed at higher levels in cluster 5 8CLCs than in 8-cell stage embryos, indicating the module is a mixture of 8-cell stage embryo and 8CLC markers. Hub gene *TENT5C* (*FAM46C*), associated with mRNA stabilization (Mroczek et al., 2017) showed upregulation in samples preceding major EGA and Yoshihara 8CLCs, hinting at its potential as a DUX4 target. *TENT5C* knockout in mice leads to male sterility, underscoring its importance in reproduction and possibly 8CLC generation. Similarly, *NEFM* expression associated with minor and major EGA stage embryos, as well as Yoshihara and Mazid intermediate and mature 8CLCs. *NEFM* is a marker of primed pluripotency (Messmer et al., 2019) and was downregulated by *LEUTX* induction in primed hPSCs (Jouhilahti et al., 2016). Expression of cell cycle-associated hub genes *CDKN1A* and *CDKN1C* varied but was upregulated in minor and major EGA embryos compared to later stages, as well as in cluster 5 and 27 8CLCs compared to hPSCs, further linking cell cycle regulation to 8CLCs, echoing findings in mouse 2CLCs. Accordingly, the 8-cell-M3 genes exhibited enrichment in GO terms associated with mesodermal differentiation and negative regulation of the mitotic cell cycle. Both unintegrated and integrative analyses suggested that 8-cell stage embryos and 8CLCs may be enriched for the G1 and G2M phases of the cell cycle. Previous studies have reported a higher proportion of cells in the G2M phase in 2CLCs compared to ESCs (Atashpaz et al., 2020; Eckersley-Maslin et al., 2016), and recent findings have highlighted a cell-cycle-associated mechanism linked to peri-nucleolar heterochromatin remodeling and *Dux* gene expression during 2CLC emergence (Zhu et al., 2021). Whether the cell cycle plays a causal role in 8CLC generation or represents a downstream effect awaits further investigation in humans, with a need to delineate between mature 8CLCs and potential intermediate states. While it is tempting to hypothesize that the cell cycle-Dux axis is conserved in humans, it is also plausible that the cell cycle may promote mechanisms beyond Dux/DUX4-induced generation of 2CLCs or 8CLCs.

To compare different 8CLC models, we integrated scRNA-seq datasets from various studies that used diverse methodologies and platforms. Despite meticulous data preprocessing and integration to minimize biases, some may remain, prompting dataset-specific analyses. Additionally, integrative analysis methods that utilize computation of meta-cells are hindered by the low absolute number of intermediate and mature 8CLCs in most datasets and therefore some results may be predominated by Yoshihara 8CLCs that are most abundant. Notably, significant differences were observed in the relative numbers of fully mature 8CLCs across datasets, impacting the applicability of these models in different experiments. Quantitatively and qualitatively, the Yoshihara and Mazid datasets stood out for containing the most representative 8-cell stage embryo-like cells. Furthermore, we observe that these 8CLCs replicate key EGA processes, such as the expression of EGA genes and TEs, and exhibit a quiescent cell state regarding energy and RNA metabolism. They also demonstrate typical EGA state regulation of alternative splicing and ribosome biogenesis, rendering them valuable models.

## ACKNOWLEDGEMENTS

We acknowledge Biomedicum Imaging Unit, University of Helsinki, with the support of the Helsinki Institute of Life Science (HiLIFE) and Biocenter Finland for imaging support, and CSC–IT Center for Science Ltd. for supercomputing. This study was supported by grants from Research Council of Finland (Academy Fellowship grants #348111 and #353549), Sigrid Jusélius Foundation, and Helsinki Institute of Life Science (HiLIFE Fellowship) to S. V.. P. P. is supported by the Finnish Cultural Foundation, Orion Research Foundation sr, Finnish Fertility Society, and the Biomedicum Helsinki Foundation. S.N. is supported by University of Helsinki Doctoral Programme in Biomedicine and the Finnish Fertility Society.

## AUTHOR CONTIRBUTIONS

P.P. and S.V. conceived the study; S.V. supervised the project; S.N. performed the immunostainings and confocal microscopy; P.P. performed bioinformatics analyses; P.P., S.N., and S.V. wrote the manuscript.

## DECLARATION OF GENERATIVE AI AND AI-ASSISTED TECHNOLOGIES IN THE WRTITING PROCESS

During the preparation of this work P.P. used ChatGPT to edit text in the manuscript. After using ChatGPT, the authors reviewed and edited the content as needed and take full responsibility for the content of the published article.

## STAR Methods

### RESOURCE AVAILABILITY

#### Lead contact

Further information and requests for resources and reagents should be directed to and will be fulfilled by the lead contact, Sanna Vuoristo.

#### Materials availability

This study did not generate new unique reagents.

#### Data and code availability

This paper analyzes existing, publicly available data. The scRNA-seq data of the 8CLC datasets were retrieved with the following accession numbers SRA: SRR14853531 (TS naïve) and SRR16975081 (TS DUX4, RNA only) (Taubenschmid-Stowers et al., 2022), SRR18215263, SRR18215264, SRR18215265, and SRR18215266 (Moya-Jodar et al., 2023), ArrayExpress: E-MTAB-10581 (Yoshihara et al., 2022), CNGB Nucleotide Sequence Archive: CNX0278328 and CNX0278329 (Mazid et al., 2022). The scRNA-seq data of naïve and primed hPSC datasets were retrieved with accession numbers SRA: SRR19353580 and SRR19353578 (Bi et al., 2022). The scRNA-seq data of human pre-implantation embryos was retrieved with accession number GEO: GSE36552 (Yan et al., 2013). The count matrix and metadata for the pre-processed human embryo datasets analyzed in Meistermann et al. (2021) were obtained from Mendeley Data repository (https://doi.org/10.17632/689pm8s7jc.1) (Radley et al., 2023). Microscopy data reported in this paper will be shared by the lead contact upon request. All original code has been deposited at Zenodo and is publicly available as of the date of publication. DOIs are listed in the key resources table. Any additional information required to reanalyze the data reported in this paper is available from the lead contact upon request.

### EXPERIMENTAL MODEL AND STUDY PARTICIPANT DETAILS

#### Naïve hESC line and culture conditions

Naive H9 (WA09, Wicell) hESCs (female), which had been previously converted from primed hESCs using the NaïveCult Induction kit (STEMCELL Technologies), were cultured in NaïveCult Expansion Medium (STEMCELL Technologies) in 5% O_2_/5% CO_2_ at 37°C. Naive hESCs were dissociated with TrypLE Express (Thermo Fisher Scientific) every 3–5 days and re-plated on mitomycin C inactivated CF1 MEF feeders (Thermo Fisher Scientific), prepared a day before hESC seeding. The cell culture medium was supplemented with 10 mM ROCKi Y-27632 (Selleckchem) for the first 24 h post plating.

### METHOD DETAILS

#### Read Alignment

The scRNA-seq raw reads were aligned to the human reference genome, GRCh38, with GENCODE (v41) transcript annotations with STARsolo (2.7.10a) (Dobin et al., 2013), using parameters -- outSAMattributes NH HI AS nM CB UB --outFilterMultimapNmax 100 --winAnchorMultimapNmax 100 --outMultimapperOrder Random --runRNGseed 777 --outSAMmultNmax 1 --soloType CB_UMI_Simple for all droplet-based (BI, TS naïve, TS DUX4, Mazid, Yoshihara, M-J) samples. The following parameters were adjusted for each dataset: --soloUMIlen 12 for Bi, TS naïve, TS DUX4, Yoshihara, and M-J; -- soloUMIlen 10 --soloCBstart 1 --soloCBlen 31 --soloUMIstart 32 for Mazid; and --soloFeatures GeneFull for TS DUX4 to adjust for introns in the nuclear RNA. The barcode whitelists were obtained from the 10X repository: 10x_v3/3M-february-2018.txt for all on 3 10x chemistry libraries (Bi, TS naive Yoshihara, and M-J) and 10x_multiome/737K-arc-v1.txt for TS_DUX4. Barcode whitelists for Mazid were received from the authors (personal communication). The Yan et al. raw reads were aligned using the following parameters --soloType SmartSeq --soloUMIdedup NoDedup --soloStrand Unstranded -- outSAMattributes NH HI AS nM --outFilterMultimapNmax 100 --winAnchorMultimapNmax 100 -- outMultimapperOrder Random --runRNGseed 777 --outSAMmultNmax 1 --limitOutSJcollapsed 3000000 --outBAMsortingBinsN 200.

#### Quality control and clustering

The filtered output from STARsolo for the scRNA-seq datasets and the Meistermann expression matrix were used for downstream processing with the R package Seurat (4.3.0). The Meistermann expression matrix was filtered to exclude the Yan et al. samples, which were pre-processed in this study. Additionally, the Fogarty OCT4 CRISPR samples were removed due to their developmental phenotype (Fogarty et al., 2017). This resulted in a matrix comprising Fogarty et al. blastocyst stage samples, Petropoulous et al. E3-E7 stage embryos, and Meistermann et al. morula and blastocyst stage samples. Cells with less than 200 RNA features and RNA features with expression in less than 3 cells were removed. Outlier cells were identified and removed based on the distribution of detected RNA features (nFeature), total counts of RNA features (nCount) and percentage of mitochondrial transcripts (percent.mt). The thresholds were adjusted for each dataset as follows: nFeature > 2500, nCount <= 30000, and percent.mt < 10 for Bi primed (5319 cells); nFeature > 2500, nCount <= 60000, and percent.mt < 10 for Bi naive (3549 cells); nFeature > 3500, nCount <= 150000, and percent.mt < 10 for TS naive (3101 cells); nFeature > 5000, nCount <= 100000, and percent.mt < 20 for TS Dux4 (1456 cells); nFeature > 1000, nCount <= 30000, and percent.mt < 10 for Mazid replicate 1 (421 cells) and 2 (359 cells); nFeature > 5000, nCount <=100000, and percent.mt < 20 for Yoshihara replicate 1 (4144 cells) and 2 (3466 cells); nFeature > 3000, nCount <= 40000, and percent.mt < 20 for MJ naive (3533 cells); percent.mt < 40 for Meistermann embryos (1597 cells). No filters were applied to Yan et al. embryo data (124 cells). The datasets were log-normalized and scaled using SCTransform to regress out percent.mt and cell cycle stage score calculated by the CellCycleScoring function. Variable features were identified with FindVariableFeatures function and PCA was performed using variable features. The number of top PCs to use in downstream analysis was determined as 30 for all datasets using the ElbowPlot function. Datasets consisting of multiple biological replicates (Mazid and Yoshihara) and the Bi naïve and primed samples were integrated (within dataset) using functions SelectIntegrationFeatures with 3000 features, PrepSCTIntegration, FindIntegrationAnchor and IntegrateData for method SCT. UMAP was implemented using top 30 PCs and cells were clustered using the Louvain algorithm and the optimized resolution parameter.

#### chooseR optimization

The chooseR algorithm (Patterson-Cross et al., 2021) was run with R (4.2) using the SCT assay, PCA reduction, 30 PCs for the dataset-wise analyses and resolutions 0.4, 0.6, 0.8, 1, 1.2, 1.4, 1.6, 1.8, 2, 4,

6, 8, 12, 16 to find optimal resolution parameter values for each dataset that were as follows: 2 for Yan, 1.8 for Bi, 1.4 for TS naïve, 1.6 for TS Dux4, 1 for Mazid, 2 for Yoshihara, 0.4 for M-J. For the integrated dataset the Harmony reduction was used with 40 PCs and the optimal resolution parameter value was determined as 2.

#### TE expression analysis

The Yan et al. data was aligned using STAR, as scTE (He et al., 2021) input requires this format for non-droplet-based methods. The alignment employed the same reference genome and GENCODE version as before but with specific parameters: --outFilterMultimapNmax 100, --winAnchorMultimapNmax 100, --outMultimapperOrder Random, --runRNGseed 777, and --outSAMmultNmax 1. For scTE, the index was constructed using the human reference genome GRCh38 and GENCODE transcript annotations (v41), along with the UCSC genome browser Repeatmasker track, using the parameter -m nointron. Droplet-based samples (BI, TS naive, TS Dux4, Mazid, Yoshihara, and M-J) were processed using scTE with the output of STARsolo and parameters -CB CB -UMI UB, while the Yan et al. samples were run with STAR output and parameters -CB False -UMI False. The scTE output matrix underwent filtering to retain only cells passing quality control in the STARsolo output and transcripts that were detected in at least three cells. The scTE expression matrix was divided into genes and TEs, treated as separate assays, and merged with the STARsolo output to create a single Seurat object for each dataset, log-normalized, and scaled after integration.

#### Fuzzy clustering

The stage-wise normalized mean expression values for TEs from Liu et al. (2019) Supplementary Data 4 was used as an input for time-course c-means Fuzzy clustering to identify embryo stage-specific expression patterns of TEs included in the Repeatmasker track used in this publication. Fuzzy clustering was performed using Mfuzz (v.2.60.0) with 12 centers and cluster membership threshold 0.6.

#### Integration of 8CLC datasets with human embryo data and clustering

The pre-processed 8CLC datasets and Yan and Meistermann oocytes and embryos were integrated using the RunHarmony function in the Harmony R package (v. 0.1.1, (Korsunsky et al., 2019)). The assay "SCT" was used, and library correspondence was provided as batch information. The chooseR algorithm was applied with top 40 PCs, using the Harmony output as reduction, and same resolutions as before. UMAP was implemented using the top 40 batch-corrected PCs, followed by clustering using the Louvain algorithm with the resolution parameter set to 2. Additionally, clustering was done with resolution parameters of 1.8 and 2.2 to visually evaluate the effect of this parameter on the formed clusters. Finally, the integrated data was log-normalized and scaled.

#### Inference of cluster identity

To evaluate the similarity of hPSCs in the 8CLC datasets to 8-cell stage embryos or to identify changes in their cellular background, we utilized the Singler R Package (1.10.0) with the Wilcox method. We employed Yan et al. oocytes and embryos, as well as Bi et al. naïve and primed hESC datasets, as references. Additionally, the AddModuleScore function from the Seurat package was used with the following gene lists to annotate the cells: Stirparo gene lists that were generated by self-organizing feature maps (SOMs) were obtained from Table S6 (Stirparo et al.), Meistermann WGCNA modules from Table S3 (Meistermann et al., 2021), Bi expression clusters from Supplementary Data 1: cluster 1 and 2, Messmer naïve and primed marker lists from Table 1, Liu et al fuzzy clustered TE lists of interest, and Göke et al. 8-cell stage specific ERV Elements from Table S1.

#### Gene regulatory network analysis

For GRN analysis, Seurat clusters with similar cell types were merged and annotated based on the results of cluster annotation and clustering with embryonic samples. We utilized the hdWGCNA (0.2.23) R package for the analysis, using groups "8-cell-like," "Yoshihara-8CLC," "Yoshihara-intermediate," "Naïve-like," and "Primed-like" as query sets. Genes expressed in at least 5% of cells in the integrated dataset were included. Metacells were constructed using the Harmony reduction and grouped based on the merged annotation and dataset information, with a minimum of 70 cells set as the threshold. An optimal soft power threshold was determined using the TestSoftPowers function and used in the network construction. Harmony batch-corrected module eigengenes were obtained, considering the origin of the datasets, and the top hub genes sorted by eigengene-based connectivity were identified for each module using the GetHubGenes function. Finally, resulting modules were manually curated to select modules specific to different 8-cell embryo like states and stem cell populations. GO enrichment analysis for “Biological process”, “Cellular component” and “Molecular function” from the GO (2023) database was done using the RunEnrichr function with all genes. The results were filtered to include only the curated modules and the adjusted p-values were calculated using Benjamini-Hochberg correction. The functions ModuleFeaturePlot and ModuleNetworkPlot, as well as the ComplexHeatmap R package (2.16.0, (Gu et al., 2016), were used for visualizing the results.

### QUANTIFICATION AND STATISTICAL ANALYSIS

#### Statistical tests and visualization

The chi-square test of independence was calculated using the chisq.test() function and the pairwise Wilcoxon rank sum tests were calculated using function pairwise.wilcox.test() in R with p-values adjusted for multiple testing with Benjamini-Hochberg correction and the upper threshold for significance set at 0.05. The following R packages were used to create visualizations of the results: ggplot2(3.4.2), viridis (0.6.2), wesanderson (0.3.6), VennDiagram (1.7.3).

#### Immunocytochemistry and imaging

Naïve H9 hPSCs were fixed on Ibidi four-well µ slides with 4% paraformaldehyde in PBS at room temperature for 15 minutes. After fixation, the cells were washed three times in washing buffer (0.1% Tween 20 in PBS) and permeabilized in 0.5% Triton X-100 in PBS at room temperature for 7 minutes, followed by a single wash. Unspecific primary antibody binding was blocked with ProteinBlock (Thermo Fisher Scientific) at room temperature for 10 minutes. Primary antibodies (rabbit anti-LEUTX, Novus Biologicals (NBP1-90890) 1/200; mouse anti-TPRX1, Merck (WH0284355M2) 1/500; rat anti-H3.X/Y, Active motif (#61161) 1/200; rabbit anti-DUX4, Abcam (ab124699) 1/300, were diluted in washing buffer and incubated at 4°C overnight. The cells were washed three times in washing buffer and incubated with secondary antibodies (anti-mouse Alexa Fluor 488 (A21202); anti-rabbit Alexa Fluor 594 (A21207); anti-rabbit Alexa Fluor 647 (A32795); anti-rat Alexa Fluor 594 (A21209), Thermo Fisher Scientific) diluted 1:1000 in washing buffer at room temperature for 25 minutes. After three washes, nuclei were counterstained with DAPI (1:1000 in washing buffer) for 10 minutes at room temperature. The cells were imaged using Leica Stellaris 8 FALCON confocal microscope with HC PL APO CS2 CORR 40×/1.25NA glycerol objective. The images were processed using Fiji (Schindelin et al., 2012).

## SUPPLEMENTAL INFORMATION AND TITLES

**Supplemental document S1. Figures S1–S5 and Tables S1-S4.**

**Supplemental Figure 1.** Supplemental information on the expression of EGA genes distinguishes between intermediate and mature 8CLC identities.

**Supplemental Figure 2.** Supplemental information on the integrated analysis of the hPSC datasets and human embryos.

**Supplemental Figure 3.** Supplemental information on intermediate 8CLCs retain stem-cell like characteristics absent in human 8-cell stage embryos.

**Supplemental Figure 4.** Supplemental information on TE expression can supplement gene expression data to distinguish between intermediate and mature 8CLCs.

**Supplemental Figure 5.** Supplemental information on GRN analysis indicates that changes in energy and RNA metabolism, alternative splicing, and ribosome biogenesis accompany the emergence of 8CLCs.

## SUPPLEMENTAL TABLES

**TABLE S1.** Basic information on hPSCs, methods used to generate or detect 8CLCs and their numbers in the original publications included in the study.

**TABLE S2.** Absolute and relative numbers of hPSCs in each 8CLC dataset annotated as embryonic stages in Yan et al. (2013) by Singler.

**TABLE S3.** Number of *TRPX1, H3Y1/2* and *LEUTX*, single, double, and triple positive cells as well as the number of *DUX4* positive cells in Yan et al. (2013) oocytes and embryos.

**TABLE S4.** Number of *TRPX1, H3Y1/2* and *LEUTX*, single, double, and triple positive cells as well as the number of *DUX4* positive cells in the 8CLC datasets.

**TABLE S5.** The number of cells from each 8CLC dataset and replicate as well as, oocytes and embryos from Yan et al. 2013 and Meistermann et al. in each integrated cluster as counts and relative frequencies of cluster total cell number and original dataset/replicate cell number.

