## Supplemental File for "Integrated transcriptomic analysis reveals a metabolically quiescent state and gene expression networks related to intermediate and mature human 8-cell stage embryo resembling cells *in vitro*"

**SUPPLEMENTAL FIGURES: TITLES AND TEXT**

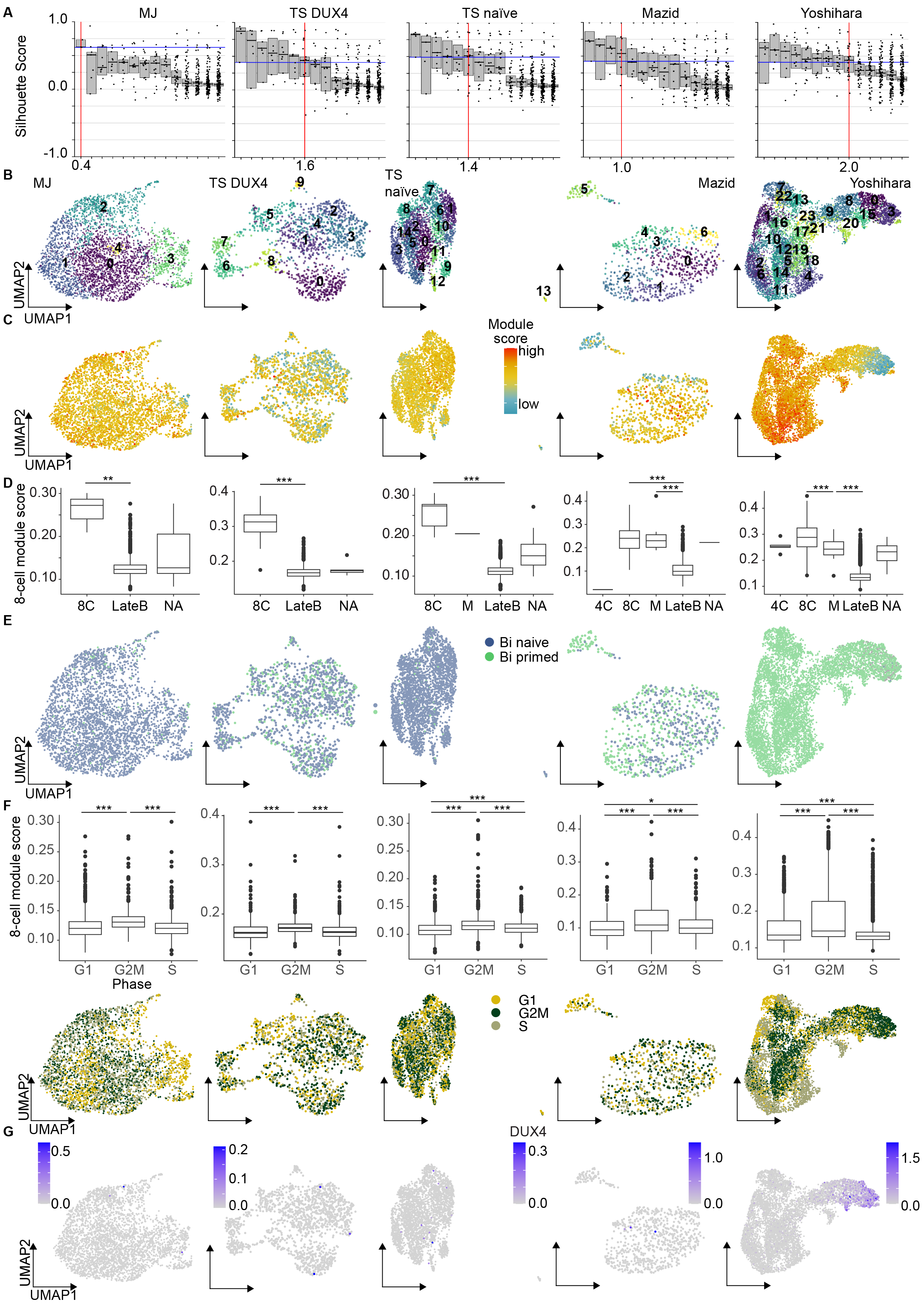

**Supplemental Figure 1. Supplemental information on the expression of EGA genes distinguishes between intermediate and mature 8CLC identities.**

A) chooseR defined silhouette scores for the tested resolution parameters to be used in Seurat clustering. The optimal parameter value is indicated by a red line. Datasets organized from left to right: MJ, Moya-Jódar; TS DUX4, Taubenschmid-Stowers DUX4-induced; TS naïve, Taubenschmid-Stowers naïve, Mazid, Yoshihara.
B) Seurat clusters as UMAPs of the hPSC datasets with cells belonging to different clusters indicated by colors.
C) Module scores for Stirparo et al. (2018) early ICM lineage-specific gene list as UMAPs of the hPSC datasets. Datasets: MJ, Moya-Jódar; TS DUX4, Taubenschmid-Stowers DUX4-induced; TS naïve, Taubenschmid-Stowers naïve.
D) Boxplots of Stirparo et al. (2018) 8-cell stage embryo-specific gene list scores in the Singler annotated cell groups for each dataset. The cells annotated as 8-cell stage embryos or morulae had highest scores. (FDR < 0.01, Wilcoxon rank sum test). Abbreviations: 4C, 4-cell stage embryo; 8C, 8-cell stage embryo; M, morula; LateB, late blastocyst; NA, undefined.
E) Module scores of the Bi et al. (2022) naïve- and primed-specific hPSC gene lists for the hPSC datasets.
F) Cell cycle phases in each dataset as an UMAP (below). The boxplot (above) shows that module scores for Stirparo et al. (2018) human 8-cell stage embryo-specific gene list were higher for cells in G2M than the other phases (FDR < 0.05, Wilcoxon rank sum test)
G) DUX4 expression in each hPSC dataset as an UMAP.

Datasets in A-G organized from left to right: MJ, Moya-Jódar; TS DUX4, Taubenschmid-Stowers DUX4-induced; TS naïve, Taubenschmid-Stowers naïve, Mazid, Yoshihara.

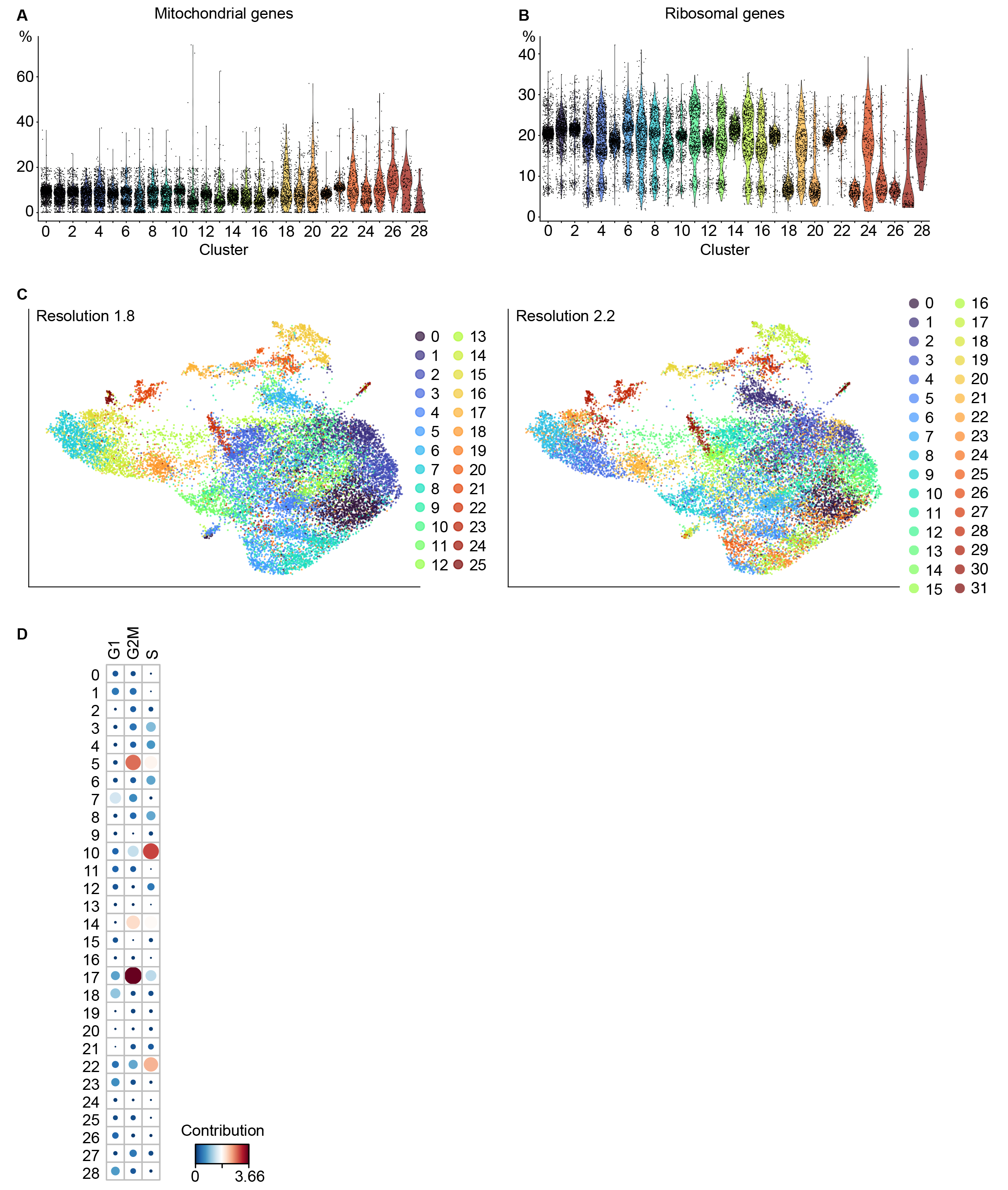

**Supplemental Figure 2. Supplemental information on the integrated analysis of the hPSC datasets and human embryos.**

A) Percentage of mitochondrial transcripts and
B) ribosomal protein-coding transcripts in each cell grouped by cluster identity. The color indicates the cluster number.
C) Seurat clustering using 20 PCs with resolution parameter set to 1.8 (left) and 2.2 (right). Cells belonging to different clusters are indicated by cluster colors.
D) The contribution in percentage of a given cell in the Chi-square matrix to the total Chi-square score calculated as: 100 % x Chi-square residuals^2 / by Chi-square statistic. The color and dot size represent the contribution in percentage.

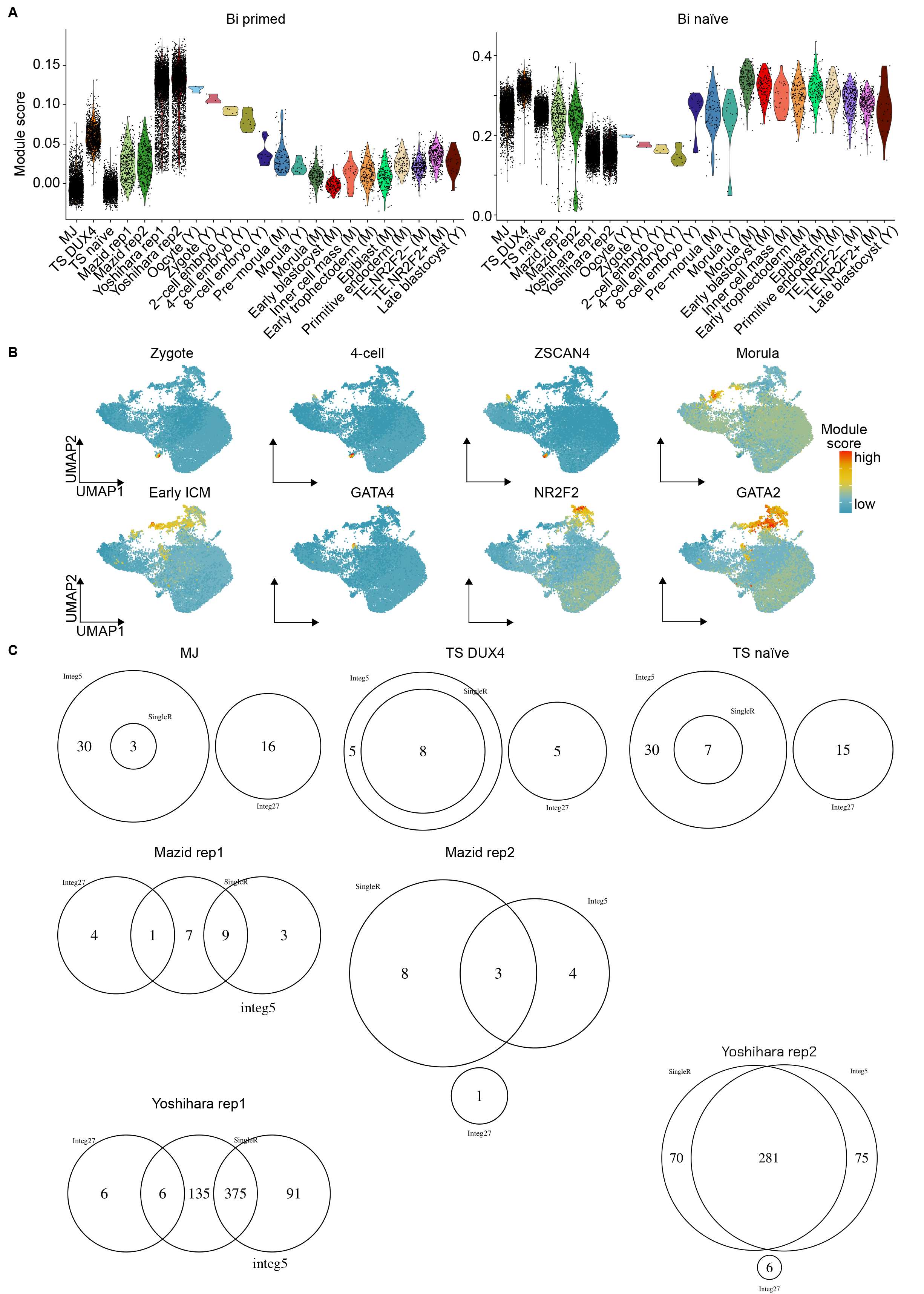

**Supplemental Figure 3. Supplemental information on intermediate 8CLCs retain stem-cell like characteristics absent in human 8-cell stage embryos.**

A) Module scores of Bi et al. (2022) naïve and primed hPSC gene lists across the original datasets and replicates, as well as the embryonic stages in Meistermann et al. (2021) and Yan et al. (2013). Datasets: MJ, Moya-Jódar; TS DUX4, Taubenschmid-Stowers DUX4-induced; TS naïve, Taubenschmid-Stowers naïve; M, Meistermann; Y, Yan. Abbreviations: rep, replicate.
B) Module scores of early embryonic (Stirparo zygote and 4-cell stage; Meistermann ZSCAN4), morula-early blastocyst stage (Stirparo compacted morula and early ICM), and lineage (Meistermann: GATA4, NR2F2, and GATA2) specific gene lists in the integrated dataset as an UMAP.
C) Venn plots of the cells annotated as 8CLCs in the dataset-wise analysis and their proportions in the 8CLC clusters 5 and 27 of the integrated dataset.

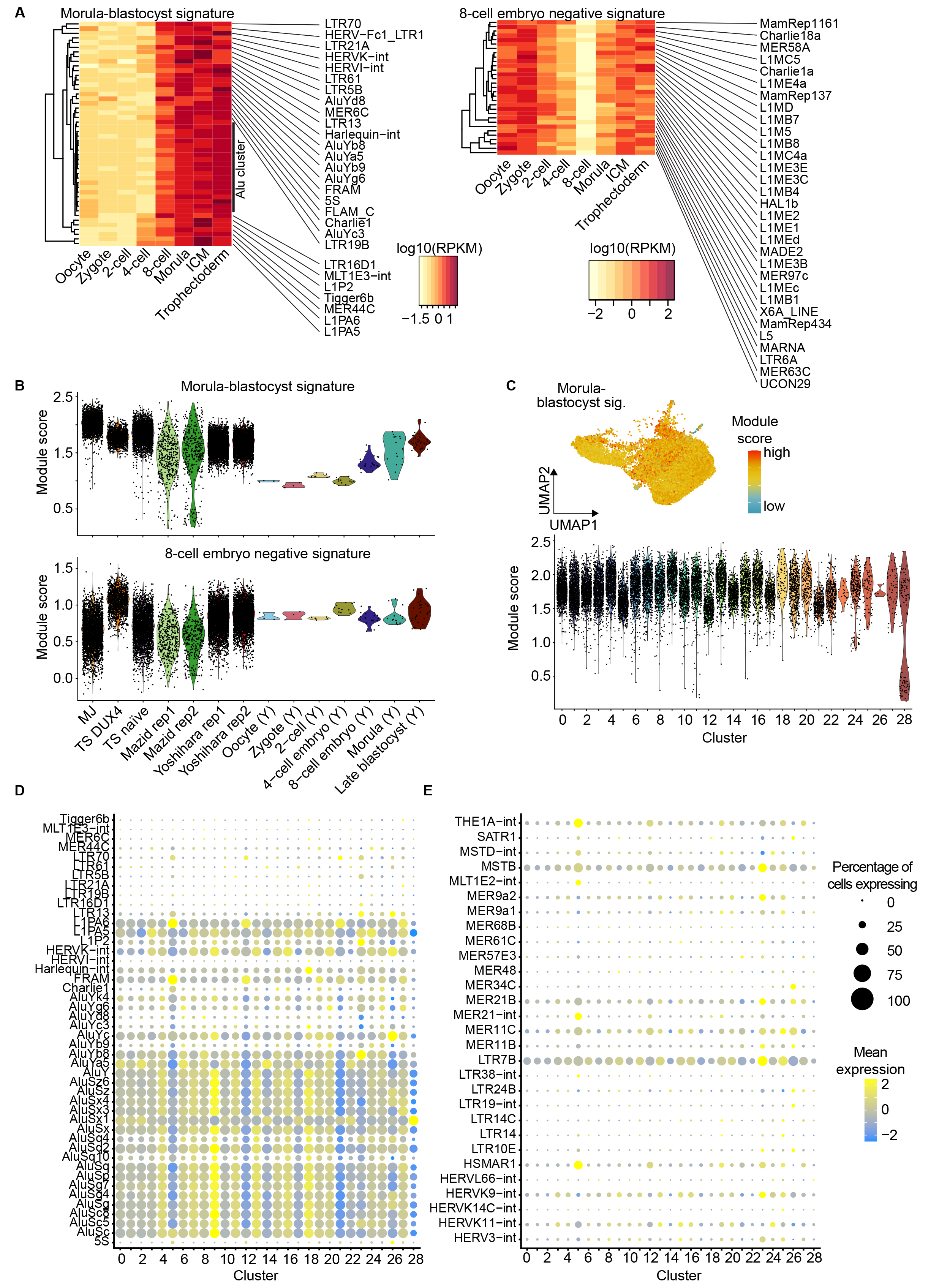

**Supplemental Figure 4. Supplemental information on TE expression can supplement gene expression data to distinguish between intermediate and mature 8CLCs.**

A) Hierarchically clustered (Euclidean distance and average linkage) heatmaps of TE expression in human oocytes and early embryos (Liu et al., 2019) for the morula-blastocyst signature (fuzzy cluster 1) and the 8-cell negative signature (fuzzy cluster 6)
B) Module scores of the morula-blastocyst and the 8-cell negative signatures in the hPSC datasets and replicates, as well as the embryonic stages in Yan et al. (2013). Datasets: MJ, Moya-Jódar; TS DUX4, Taubenschmid-Stowers DUX4-induced; TS naïve, Taubenschmid-Stowers naïve; Y, Yan. Abbreviations: rep, replicate.
C) Module scores of the morula-blastocyst signature in the integrated dataset as an UMAP (top) and across the clusters (bottom).
D) Dotplot of TE expression across clusters in the integrated dataset with cluster average expression (scaled) indicated in color and percentage of cells expressing TE as dot size for the morula-blastocyst and
E) the 8-cell-morula signatures.

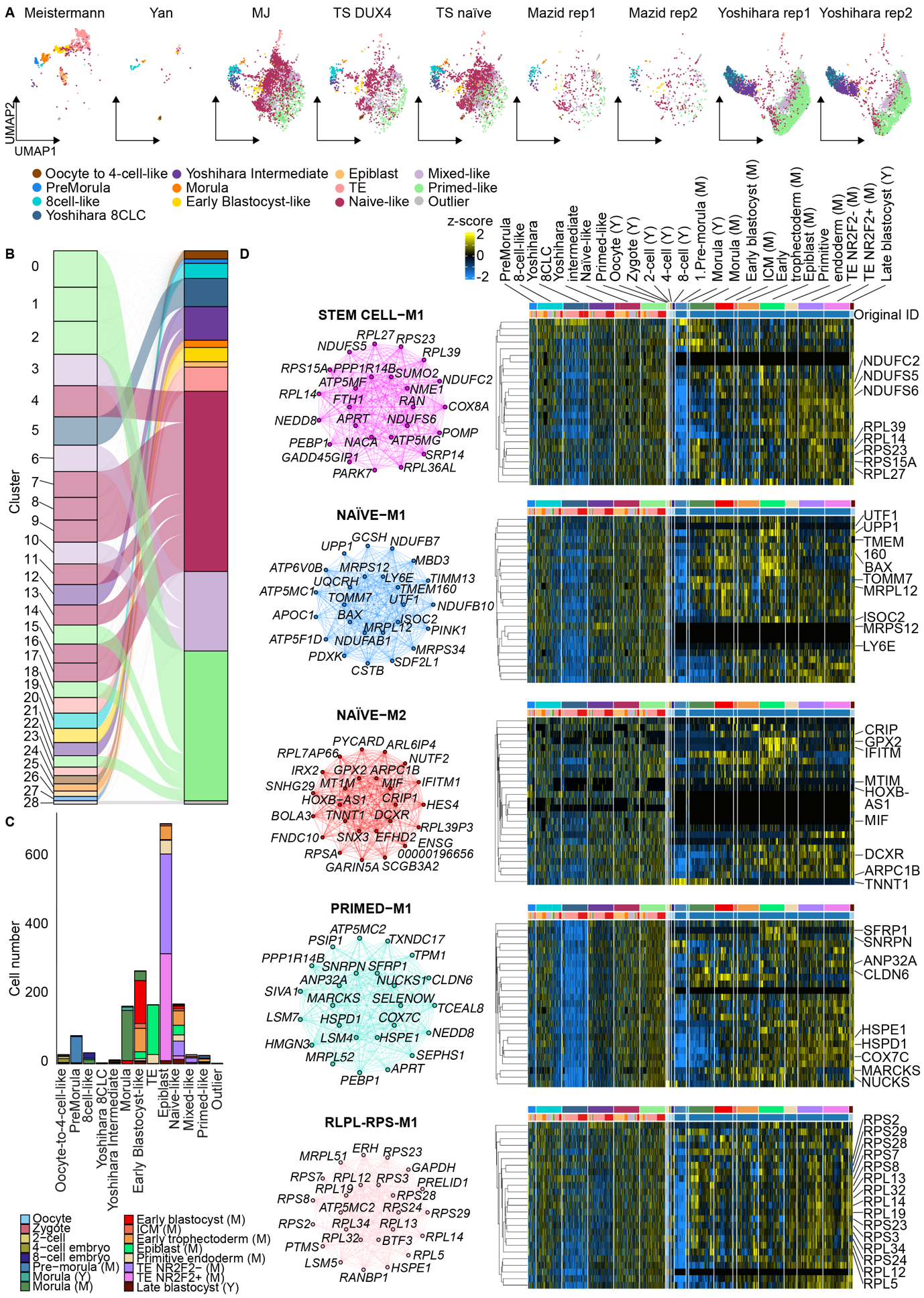

**Supplemental Figure 5. Supplemental information on GRN analysis indicates that changes in energy and RNA metabolism, alternative splicing, and ribosome biogenesis accompany the emergence of 8CLCs.**

A) UMAP of the integrated data split by original dataset and replicate with the cell groups used for GRN analysis indicated in colors. Datasets: MJ, Moya-Jódar; TS DUX4, Taubenschmid-Stowers DUX4-induced; TS naïve, Taubenschmid-Stowers naïve; M, Meistermann; Y, Yan. Abbreviations: rep, replicate.  Clusters have been combined to bigger entities based on similarity as depicted in B).
B) with an alluvial plot showing the correspondence between cluster (left) and GRN cell grouping (right) indicated by the color key in A).
C) The number of embryonic samples (Meistermann et al., 2021; Yan et al., 2013) in the groups used as input for the GRN analysis. The colors indicate the embryonic stage or lineage.
D) Network plots (left) and hierarchically clustered (Euclidean distance and average linkage) expression heatmaps (right) of hub genes for GRNs representative of stem cell-specific states. Heatmap colors indicate z-scores that have been calculated in a dataset- and replicate-wise manner. Abbreviations: ICM, inner cell mass; TE, trophectoderm; M, Meistermann et al.; Y, Yan et al.

**SUPPLEMENTAL TABLES**

**Table S1.** Basic information on hPSCs, methods used to generate or detect 8CLCs and their numbers in the original publications included in the study.

| **Dataset** | **Cell type** | **Optimal condition for 8CLC generation in original publication** | **Percentage of 8CLCs in original publication** | **8CLC identification method** |
| --- | --- | --- | --- | --- |
| Mazid  (Mazid et al., 2022) | primed-to-naive hPSCs | e4CL-D5 | 11.90 % | clustering |
| MJ  (Moya-Jódar et al., 2023) | naive human induced PSCs | NA | ~0.3% | ZSCAN4+/ MLT2A1+ expression |
| TS DUX4  (Taubenschmid-Stowers et al., 2022) | naive hESCs | 2-4h DUX4 induction | not reported | clustering |
| TS naïve  (Taubenschmid-Stowers et al., 2022) | naive hESCs | NA | 1.61 % | clustering |
| Yoshihara  (Yoshihara et al., 2022) | primed hESCs | 15-minute DUX4 pulse followed by a 12-hour incubation | 6.60 % | SingleR with the scRNA-seq data of preimplantation embryos and hESCs (Yan et al., 2013) as reference |

**Table S2.** Absolute and relative numbers of hPSCs in each 8CLC dataset annotated as embryonic stages in Yan et al. (2013) by Singler.

|  | **Counts** | | | |
| --- | --- | --- | --- | --- |
| **Dataset** | 4-cell | 8-cell | Morulae | Late blastocyst |
| Mazid | 1 | 28 | 8 | 742 |
| MJ | 0 | 3 | 0 | 3503 |
| TS DUX4 | 0 | 8 | 0 | 1443 |
| TS naive | 0 | 7 | 1 | 3072 |
| Yoshihara | 5 | 867 | 29 | 6638 |
|  | **Percentages** | | | |
| **Dataset** | 4-cell | 8-cell | Morulae | Late blastocyst |
| Mazid | 0.13 | 3.59 | 1.03 | 95.13 |
| MJ | 0 | 0.08 | 0 | 99.15 |
| TS Dux4 | 0 | 0.55 | 0 | 99.11 |
| TS naive | 0 | 0.23 | 0.03 | 99.06 |
| Yoshihara | 0.07 | 11.39 | 0.38 | 87.23 |

**Table S3.** Number of *TRPX1, H3Y1/2* and *LEUTX*, single, double, and triple positive cells as well as the number of *DUX4* positive cells in Yan et al. (2013) oocytes and embryos.

|  | **Counts** | | | | |
| --- | --- | --- | --- | --- | --- |
| **Stage (Yan et al., 2013)** | **TPRX1_LEUTX_H3Y** | **TPRX1_LEUTX** | **TPRX1_H3Y** | **LEUTX_H3Y** | **DUX4** |
| Oocyte | 0 | 0 | 0 | 0 | 0 |
| Zygote | 0 | 0 | 0 | 0 | 1 |
| 2-cell | 1 | 2 | 1 | 2 | 0 |
| 4-cell | 3 | 3 | 3 | 12 | 3 |
| 8-cell | 20 | 20 | 20 | 20 | 3 |
| Morulae | 15 | 15 | 16 | 15 | 1 |
| Late blastocyst | 0 | 0 | 0 | 0 | 0 |
|  | **Percentages** | | | | |
| **Stage (Yan et al., 2013)** | **TPRX1_LEUTX_H3Y** | **TPRX1_LEUTX** | **TPRX1_H3Y** | **LEUTX_H3Y** | **DUX4** |
| Oocyte | 0 | 0 | 0 | 0 | 0 |
| Zygote | 0 | 0 | 0 | 0 | 33.33 |
| 2-cell | 16.67 | 33.33 | 16.67 | 33.33 | 0 |
| 4-cell | 25 | 25 | 25 | 100 | 25 |
| 8-cell | 100 | 100 | 100 | 100 | 15 |
| Morulae | 93.75 | 93.75 | 100 | 93.75 | 6.25 |
| Late blastocyst | 0 | 0 | 0 | 0 | 0 |

**Table S4.** Number of *TRPX1, H3Y1/2* and *LEUTX*, single, double, and triple positive cells as well as the number of *DUX4* positive cells in the 8CLC datasets.

|  | **Counts** | | | | |
| --- | --- | --- | --- | --- | --- |
| **Dataset** | **TPRX1_LEUTX_H3Y** | **TPRX1_LEUTX** | **TPRX1_H3Y** | **LEUTX_H3Y** | **DUX4** |
| Mazid | 9 | 12 | 88 | 10 | 0 |
| MJ | 6 | 7 | 17 | 11 | 3 |
| TS DUX4 | 5 | 5 | 7 | 15 | 3 |
| TS naive | 13 | 13 | 18 | 22 | 0 |
| Yoshihara | 3771 | 3772 | 3925 | 7134 | 1048 |
|  | **Percentages** | | | | |
| **Dataset** | **TPRX1_LEUTX_H3Y** | **TPRX1_LEUTX** | **TPRX1_H3Y** | **LEUTX_H3Y** | **DUX4** |
| Mazid | 1.15 | 1.54 | 11.28 | 1.28 | 0 |
| MJ | 0.17 | 0.2 | 0.48 | 0.31 | 0.08 |
| TS DUX4 | 0.34 | 0.34 | 0.48 | 1.03 | 0.21 |
| TS naive | 0.42 | 0.42 | 0.58 | 0.71 | 0 |
| Yoshihara | 49.55 | 49.57 | 51.58 | 93.75 | 13.77 |
